# Neuronal cell-type and circuit basis for visual predictive processing

**DOI:** 10.1101/2021.12.31.474667

**Authors:** Jinmao Zou, Lawrence Huang, Lizhao Wang, Yuanyuan Xu, Chenchang Li, Qilin Peng, Hongkui Zeng, Siyu Zhang, Lu Li

## Abstract

Bayesian Brain theory suggests brain utilises predictive processing framework to interpret the noisy world^1-11^. Predictive processing is essential to perception, action, cognition and psychiatric disease^12^, but underlying neural circuit mechanisms remain undelineated. Here we show the neuronal cell-type and circuit basis for visual predictive processing in awake, head-fixed mice during self-initiated running. Preceding running, vasoactive intestinal peptide (VIP)-expressing inhibitory interneurons (INs) in primary visual cortex (V1) are robustly activated in absence of structured visual stimuli. This pre-running activation is mediated via distal top-down projections from frontal, parietal and retrosplenial areas known for motion planning, but not local excitatory inputs associated with the bottom-up pathway. Somatostatin (SST) INs show pre-running suppression and post-running activation, indicating a VIP-SST motif. Differential VIP-SST peri-running dynamics anisotropically suppress neighbouring pyramidal (Pyr) neurons, preadapting Pyr neurons to the incoming running. Our data delineate key neuron types and circuit elements of predictive processing brain employs in action and perception.

**One Sentence Summary:** VIP-SST-Pyr microcircuit in visual predictive processing during voluntary running

## Main

To make probabilistic inference of the changing environment, brain may employ a hierarchical generative framework called predictive processing^1-11^. In this framework, an internal model (the upper layer) generates an ‘initial guess’ or prediction; the lower layer reports up the prediction-error, which is the discrepancy between the prediction and reality so the internal model can make a better guess. By repetitions, predictive processing minimises the prediction-error, leading to successful inference of the real cause so that perception or cognition may arise. From Bayesian perspectives, it lays a foundation for unifying perception, action and cognition, and accordingly, less optimized predictive processing may account for psychiatric disorders^12^. However, how the brain neurobiologically implements such a mathematically convincing framework remains largely unknown.

In motor tasks, frontal, parietal and retrosplenial brain regions fire preparatory neural activities preceding the action initiation, which could be neural correlates of predictive processing^13-20^. Recent studies have reported prediction-error encoding neurons in visual, auditory and somatosensory cortices^21-24^, but understandings of the circuit mechanisms remain elusive. Firstly, neural computations are carried out by brain circuits consisting of interconnected neuronal cell-types, i.e. distinct groups of neurons categorised by characteristic morphology, physiology, connectivity and genetics^25-28^, however, key neuron types in predictive processing are undefined. Secondly, predictive processing, by its mathematical definition, requires interplay between bottom-up and top-down processes^1,8,11,29,30^, yet underlying circuit elements are less delineated. Finally, predictive processing requires neuronal dynamics of both excitatory pyramidal (Pyr) cells and INs, but functional data of neuron populations that elucidate neuronal activity dynamics at single-neuron resolution are largely absent.

We investigated the neuronal cell-type and circuit basis for predictive processing in awake, head-fixed mice during brief, sporadic self-initiated running (Fig. 1a and Supplementary Fig. 1, also see Methods). Self-initiated actions employ predictive processing to suppress self-induced sensory outcomes^2,4,8^ and are essential to human perceptual, cognitive and social experiences, making it an ideal behavioural paradigm to dissect neural circuits of predictive processing in experimental animals. Without oversimplification, we propose that internal models in cortical motion-planning centres predict visual outcomes of running; the predictions, likely encoded by preparatory neural activities in frontal, parietal or retrosplenial brain regions^13-20^, are sent to V1 via top-down projections to suppress running-induced visual responses^31^ thus result in observable pre-running neuronal signature in V1; prediction-error encoding neurons in V1 then sent back prediction-errors through reciprocal connections between V1 and motion-planning areas^23,31^. Based on this working model, we utilised transgenic mouse lines^32,33^ that cell-type specifically expressed genetically encoded Calcium (Ca) indicators^34^ (GECIs, e.g. GCaMP6s) to identify key neuron types in predictive processing. To reveal the pre-running neuronal signature, we performed *in vivo* two-photon (2-p) Ca imaging of V1 layer (L) 2/3 genetically-defined neuron populations^35^.

**Figure 1.**
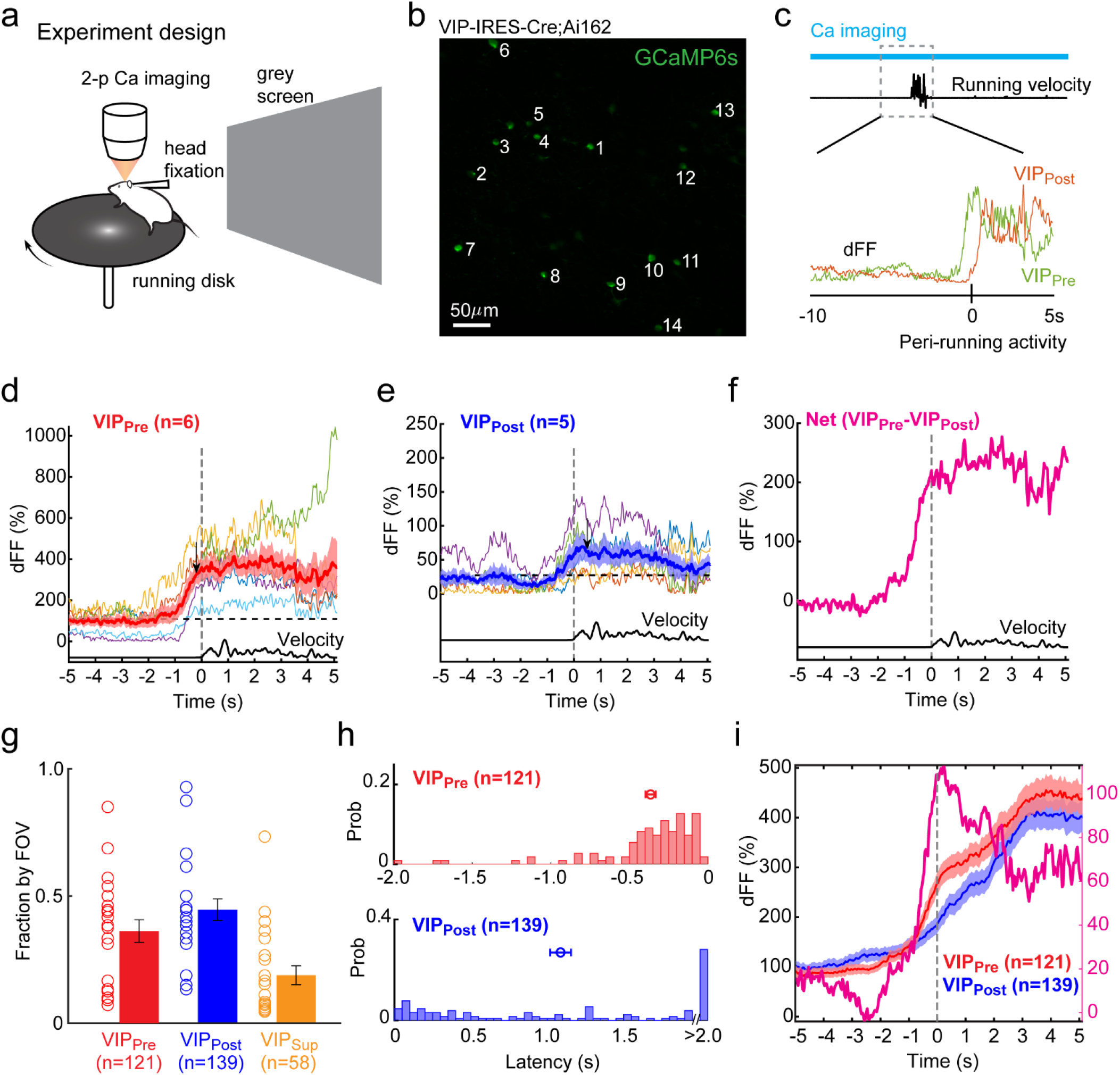
Robust activation of V1 VIP neurons prior to self-initiated running. **a**. Experimental setup showing Ca Imaging in naïve head-fixed mice during self-initiated running in absence of structured visual stimulations. **b**. Example field of view (FOV, imaged at 174 µm sub-pial) in a VIP-Cre;Ai162 (GCaMP6s) mouse. Numbers on the Z-projection (time series) image of the FOV indicate regions of interest (ROIs). **c**. Peri-running analysis of ΔF/F by Peri-Running Time Histograms (PRTHs). Bottom shows example VIP_Pre_ and VIP_Post_ neurons identified. **d – f**. Time courses of peri-running activities of VIP_Pre_ (**d**), VIP_Post_ neurons (**e**) in **b** and ‘Net’ activation (**f**). Thin and thick solid lines plot individual cells and group averages, respectively. Bottom trace plots running velocity. Arrows mark mean latency. Note the substantial activity increase within ∼1 s prior to running initiation. For display purposes, only ΔF/F within -5 sec to 5 sec from the running onset is shown. Horizontal dashed line: pre-running baseline; vertical dashed line: running onset; Shade area: s.e.m. **g – i**. Population data showing fractions of VIP_Pre_, VIP_Post_ and VIP_Sup_ neurons by FOV (**g**); latency distributions of VIP_Pre_ and VIP_Post_ neurons (**h**); average time courses of VIP_Pre_, VIP_Post_ populations and net pre-running activation (**i**). Error bars: s.e.m.

We chose to first focus on V1 L2/3 VIP INs due to their extensive innervations by top-down projections^36^ and importance demonstrated in predictive processing-associated cognitive functions such as learning and attention^37-41^.

### VIP INs are robustly activated *prior to* self-initiated running in absence of structured visual stimuli

We imaged 335 L2/3 VIP INs with 400 × 400 µm fields of view (FOV, n = 22, Fig. 1b) in naïve VIP-Cre;Ai162 (GCaMP6s) mice. Mice were free to run at arbitrary times relative to the imaging start during 200 sec imaging sessions^42^ (Supplementary Fig. 1). To identify neural correlates of predictive processing, we quantified peri-running (−10 to 5 sec from running onset, Fig. 1c) fluorescence changes (ΔF/F) and constructed Peri-Running Time Histograms (PRTHs) for individual fluorescently active regions of interest (ROIs, n = 318). As shown by the example FOV in Fig. 1b, VIP INs significantly increases ΔF/F during running (Supplementary Fig. 2), consistent with the activity augmentation reported during locomotion^42^. However, 46% (6/13) of active ROIs significantly increases ΔF/F *before* the running onset (latency: -0.192 ± 0.043 sec, mean ± s.e.m, n = 6, Fig. 1d), whereas in the rest 38% (5/13) of ROIs, significant ΔF/F increase occurs after running starts (latency: 0.486 ± 0.381 sec, n = 5, Fig. 1e & Supplementary Fig. 2). For convenience, these neurons are termed VIP_Pre_ and VIP_Post_, respectively. At population level, VIP_Pre_ INs (36.2 ± 4.3%, total: 121/318) distribute extensively across all 22 FOVs (Fig. 1g) with an average latency at -0.365 ± 0.033 sec (n = 121, Fig. 1h). The fast rise of pre-running activity in mean PRTH (Fig. 1i) suggests VIP_Pre_ receive strong and synchronous excitatory inputs prior to running initiation, e.g. the top-down preparatory neural activities. VIP_Post_ INs, which consist of 44.6 ± 4.1% (total: 139/318) of the entire VIP population, show an average latency of 1.087 ± 0.068 sec (n = 139) with a slow up-ramping PRTH (Fig. 1g-i). In the rest 17.1 ± 4.1% (n = 58/318) of VIP INs, activities are slightly suppressed (Supplementary Fig. 3). Consistent with previous studies^42,43^, both VIP_Pre_ and VIP_Post_ show similar sustained activity elevation after running start (Supplementary Figs. 3 – 5), suggesting common sets of excitatory sources responsible for post-running activation in both VIP_Pre_ and VIP_Post_ INs. This allows us to estimate the ‘net’ pre-running VIP activation by subtracting the average activity of VIP_Post_ from that of VIP_Pre_ INs, which reveals a sharp, transient pre-running VIP activity (Fig. 1i) mimicking preparatory neural activities (e.g. the Readiness Potential) reported in human, monkeys and rodents^13-20^. Pre-running activation of VIP INs is NOT an evoked response to uncontrolled visual input, because robust, even slightly stronger pre-running VIP activation is found in mice running in darkness (112 ROIs, 6 FOVs in 3 VIP-Cre;Ai162 mice, Supplementary Figs. 4&5). Different temporal activations of co-existing VIP_Pre_ and VIP_Post_ INs differentiate our findings from continuous locomotion, in which activity augments are global, non-specific and depend on neuromodulator-mediated arousal/brain state changes^42,43^.

In our behavioural paradigm, running occurs randomly (Supplementary Fig. 1). This randomness validates running is self-initiated thus adequately recruits predictive processing, but makes it infeasible to deliver manipulations prior to running because now when the animal will run is unknown to the experimenter. As we show below, pre-running activation is cell-type specific to VIP INs, ruling out possibilities of artifacts by undocumented movement and/or data processing/analysis. More importantly, on average 5.5 ± 0.85 VIP_Pre_ INs are activated prior to running, and 68% (15/22) of FOVs have at least 4 VIP_Pre_ INs identified (Supplementary Fig. 3). This clustering of VIP_Pre_ strongly suggests common causes for VIP pre-running activation instead of stochastic fluctuations of individual VIP INs. In summary, our data demonstrate an early engagement of L2/3 VIP INs preceding self-initiated running, providing direct *in vivo* evidence that the VIP network plays crucial roles in visual predictive processing.

### Distal top-down projections provide dominant excitatory drive to pre-running VIP activation

V1 VIP INs receive top-down, bottom-up and local recurrent excitatory inputs^36-40^. To delineate the interplay between top-down and bottom-up pathways pertaining to pre-running VIP activation, we examined the peri-running activities of V1 local L2/3 Pyr neurons (n = 378 ROIs in 9 FOVs, Fig. 2 & Supplementary Figs. 6&7) in Emx1-IRES-Cre;CaMK2a-tTA;Ai94 mice (expressing GCaMP6s in pan-excitatory cortical neurons). Here L2/3 Pyr activities act as the readout of bottom-up and local excitatory influences on VIP INs, because LGN and L4 bottom-up pathways provide the driving input to L2/3 Pyr neurons. Furthermore, if pre-running VIP activation is indeed part of visual predictive processing, characteristic neural dynamics, especially Pyr activity suppression, should be observed as predicted by our working model. Our data confirm this projection. As exemplified in Fig. 2a-d, contrary to VIP, only a tiny portion of Pyr neurons (2%, 1/47 ROIs) are Pyr_Pre_ cells (Fig. 2b); 43% (20/47) and 21% (10/47) are Pyr_Post_ cells (Fig. 2c) and non-responsive (Supplementary Fig. 6), respectively. Thirty-four percent (16/47) of Pyr neurons, as expected, show pre-running suppression (Pyr_Sup_, Fig. 2d). Across all three mice with running behaviour similar to VIP animals (Supplementary Fig. 7), Pyr_Pre_ cells are consistently scarce (1.9 ± 0.8%, median: 0, total: 8/378, Fig. 2e&f), indicating the pre-running activation is an intrinsic property of VIP INs. The scarcity of Pyr_Pre_ neurons suggests bottom-up and local pathways provided little, if any, excitatory drive to pre-running VIP activation. Instead, the fact that the majority of Pyr neurons (64.9 ± 6.8%, median: 66.7%, total: 252/378) are Pyr_Post_ cells indicate local Pyr activities contribute more to post-running VIP activation. These results are consistent with previous studies showing LGN and L4 only slightly increase activity during, but not before locomotion^43,44^. Importantly, in 30.9 ± 6.6% (median: 32.0%, total: 118/378) of Pyr neurons, we find profound pre-running suppression (Fig. 2h & Supplementary Fig. 7), providing experimental support that the observed peri-running dynamics of Pyr neurons are recruited by visual predictive processing in voluntary running. Taken together, these data demonstrate bottom-up/local excitatory inputs are insufficient to drive the pre-running activation of VIP INs.

**Figure 2.**
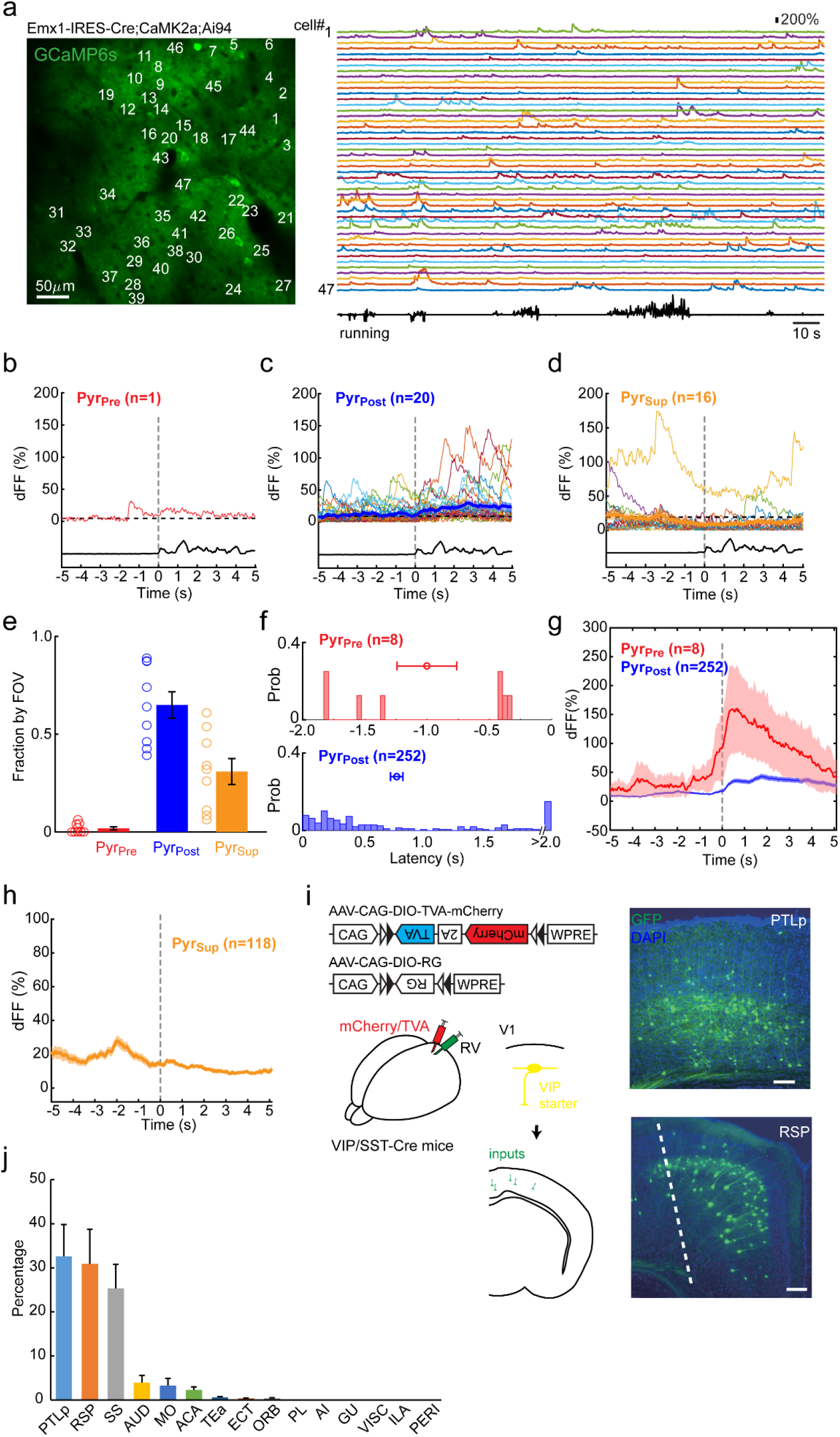
Distal top-down, but not local excitatory inputs, drive pre-running activation of VIPPre neurons. **a**. Example FOV from an Emx1-IRES-Cre;CaMKIIa-tTA;Ai94 (GCaMP6s) mouse. Data display is similar to Fig. 1. **b** – **d**. Peri-running analysis of pyramidal cells in **a**. Activities of Pyr_Pre_ (**b**), Pyr_Post_ (**c**) and Pyr_Sup_ cells (**d**) are plotted. Note only 1 (2%) Pyr_Pre_ neuron among 47 Pyr cells imaged. **e - h**. Population data showing proportions of Pyr_Pre_, Pyr_Post_ and Pyr_Sup_ neurons per FOV (**e**); latency distributions of Pyr_Pre_ (mean ± SEM: -1.000 ± 0.240 s) and Pyr_Post_ (0.758 ± 0.046 s) cells (**f**); average time courses of Pyr_Pre_, Pyr_Post_ (**g**) and Pyr_Sup_ populations (**h**). The scarcity of Pyr_Pre_ neurons across all FOVs, which proves the pre-running activation of VIP INs is not running or data analysis artifacts, strongly suggests local inputs are insufficient to drive pre-running VIP activation. **i&j**. VIP INs receive direct mono-synaptic top-down excitatory projections from frontal, parietal and retrosplenial motion-planning brain regions. **i**. Left: Schema of retrograde tracing of pre-synaptic inputs of V1 VIP neurons with engineered Cre-dependent rabies virus. Right: Example coronal section images showing dense fluorescently labeled input neurons in lateral posterior parietal cortex (PTLp, top) and retrosplenial cortex (RSP, bottom), respectively. Scale bar: 100µm. **j**. Quantification of neocortical monosynaptic inputs to V1 L2/3 VIP neurons by cortical regions. Extrastriatal inputs to VIP neurons are dominated by PTLp, RSP and Somatosensory (SS) cortices, which are known to fire preparatory activities before voluntary actions. Brain regions are ranked by their relative input strength, from high to low. AUD: Auditory cortex; MO: Motor cortex; ACA: Anterior Cingulate cortex; TEa: Temporal association areas; ECT: Ectorhinal area; ORB: Orbital area; PL: Prelimbic area; AI: Agranular insular area; GU: Gustatory areas; VISC: Visceral area; ILA: Infralimbic area; PERI: Perirhinal area. Error bars: s.e.m.

To further determine the excitatory source of pre-running VIP activation, we conducted Cre-dependent retrograde mono-synaptic tracing of distal inputs to V1 VIP INs with engineered rabies virus in VIP-IRES-Cre;Ai14 mice. Unilateral viral injections were made at superficial locations of V1, and we quantified ipsilateral neocortical input density to V1 VIP INs. Our tracing results confirm V1 L2/3 VIP INs are directly innervated by distal monosynaptic excitatory top-down projections from ipsilateral motion-planning cortical regions^36^ (Fig. 2i&j). Fluorescent positive presynaptic cells had typical morphology of projection neurons and distribute mostly in deep cortical layers (Fig. 2i). Areal distribution is widespread but biased, with more than 85% inputs originated from lateral posterior parietal cortex (PTLp), retrosplenial (RP), and somatosensory (SS) cortices (Fig. 2j), which are cortical areas known responsible for motion-related preparatory neural activities in rodents^13-20^. These monosynaptic excitatory top-down projections, which convey predictions by the ‘upper layer’, provide the anatomical substrate required by the fast, time-locked pre-running activation of VIP INs we observe in V1.

### VIP-SST motif is essential to visual predicative processing

In mouse V1, INs consist of three complete, non-overlapping types, namely Parvalbumin (PV)-, SST- and 5HT3aR-expressing INs^45^ (VIP is a main sub-group of 5HT3a+ INs). SST INs are responsible for feedback inhibition in bottom-up information processing and important to learning, attention and orientation selectivity^37-42Error!Bookmark not defined.^. Meanwhile, SST INs are the primary post-synaptic target of neighboring VIP INs^37-42,46^, and we have previously discovered that two subpopulations of V1 L2/3 SST INs are closely associated with the bottom-up and top-down pathway, respectively^35^. These results indicate SST INs may play crucial roles in top-down and bottom-up crosstalk of predictive processing. To further dissect circuit mechanisms for visual predictive processing, we imaged and analyzed peri-running activities of L2/3 SST INs (n = 304 ROIs in 23 FOVs) using SST-IRES-Cre;Ai163 (GCaMP6s) mice. Consistent with published data^35,41,42^, SST INs are highly spontaneously active in awake V1 (Fig. 3a). However, unlike VIP INs, much fewer SST INs are activated prior to running initiation (Fig. 3b & c). On average only 5.8 ± 1.9% of SST INs (median: 0, total: 17/284) are SST_Pre_ cells (Fig. 3d – f, Supplementary Figs. 8&9), with all showing post-running activity overwhelms that of pre-running (Fig. 3f). This *de facto* weak pre-running SST activity, which is in sharp contrast with VIP INs, dismisses VIP_Pre_ pre-running activation is caused by coincidental spontaneous firing or undocumented movement. It further suggests preparatory neural activities via the top-down pathway have relatively little contribution to overall SST activation. About 67.1 ± 4.7% (median: 63.6%, total: 186/284) and 27.1 ± 4.6% (median: 25%, total: 81/284) of SST INs show post-running activation (SST_Post_) and pre-running suppression (SST_Sup_, Fig. 3d – g, Supplementary Fig. 9), respectively. The substantial pre-running SST suppression confirms SST INs are the downstream target of pre-running activated VIP^46^ and supports a novel VIP-SST motif in visual predictive processing. This finding is consistent with previous studies of anatomical connectivity^46^ and predictive processing-associated cognitive functions such as attention and learning^37-41^, thus represents a circuit motif different from the SST-VIP circuit reported for the plasticity of visual mismatch signal (i.e. prediction-error) processing during development^47^. In our data, more than half of SST INs are SST_Post_ cells, which are closely associated with bottom-up and local excitatory pathways, highly consistent with our previous finding of two SST subtypes^35^. Functionally, pre-running SST suppression induces early disinhibition, and post-running SST activation causes delayed inhibition to neighbouring postsynaptic Pyr neurons, respectively. This spatiotemporally coordinated activation of the VIP-SST motif may result in a synergistic ‘push-pull’ like regulation of the responsiveness of Pyr neurons. To summarise, our results demonstrate the VIP-SST motif is a key element of neural circuits for predictive processing.

**Figure 3.**
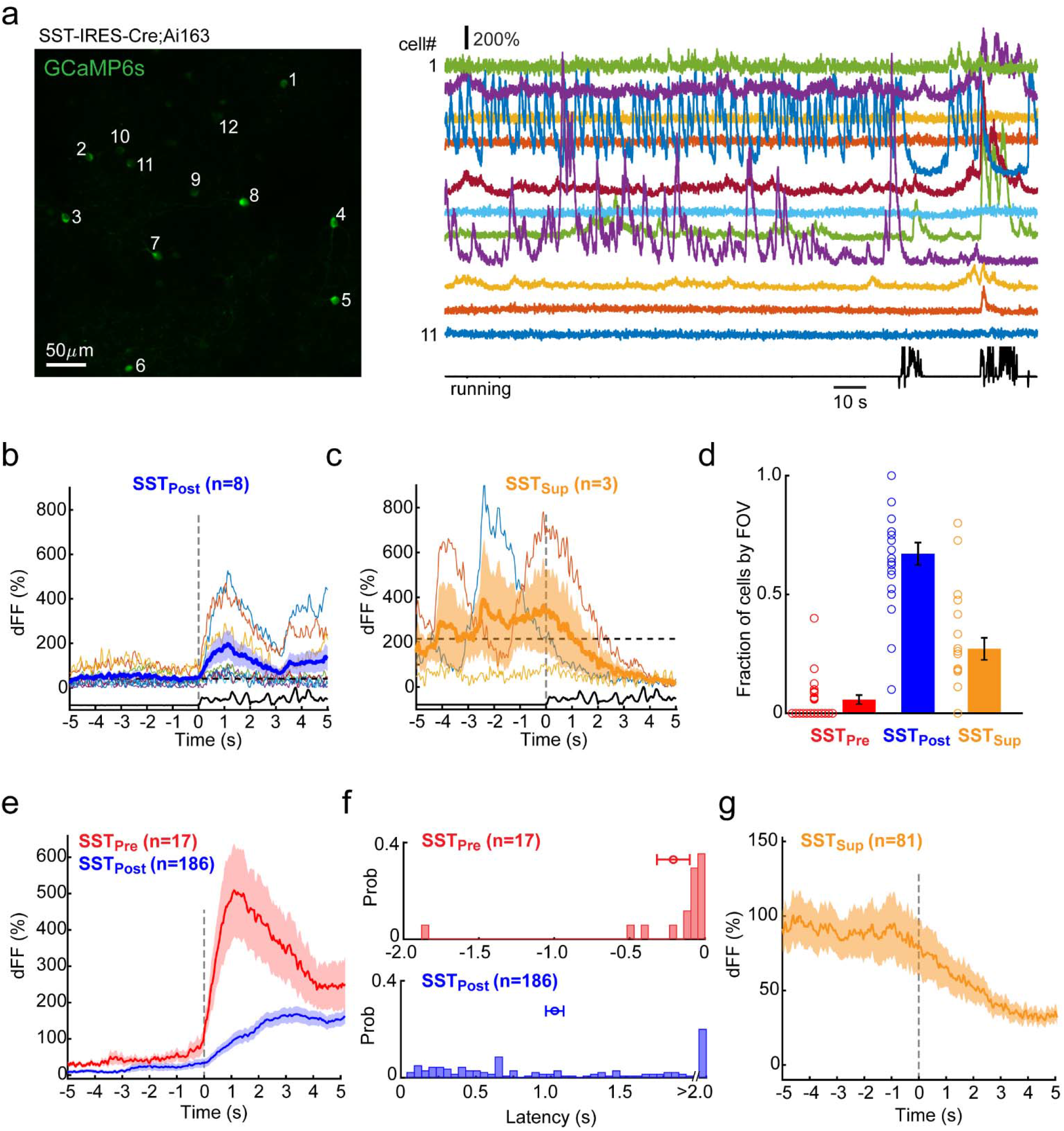
VIP-SST motif is a key circuit component of visual predictive processing. **a**. Example FOV in an SST-Cre;Ai163 (GCaMP6s) mouse. **b – d**. Peri-running analysis of SST neurons in **a**. Activities of 8 SST_Post_ (**b**) and 3 SST_Sup_ cells (**c**) are plotted. Note this FOV has no SST_Pre_ cells despite high spontaneous activity, indicating VIP_Pre_ is NOT caused by coincidental spontaneous firing or undocumented movement. **e – g**. Population data showing profound pre-running suppression and post-running activation in SST neurons. Individual panels show fractions of SST_Pre_, SST_Post_ and SST_Sup_ neurons by FOV (**e**); latency distributions of SST_Pre_ (mean ± s.e.m: -1.000 ± 0.240 sec), SST_Post_ (0.758 ± 0.046 sec) cells, respectively (**f**); average time courses of Pyr_Pre_, Pyr_Post_ (**g**) and Pyr_Sup_ populations (**h**). Note that high spontaneous firing does not necessarily result in robust and strong pre-running activation like that in VIP INs. Furthermore, unlike VIP_Pre_, SST_Pre_ neurons typically have strong post-running activation, which overwhelms the pre-running activities. Error bars: s.e.m.

### Spatiotemporally coordinated peri-running dynamics in the VIP-SST-Pyr microcircuit adaptively regulate V1 visual information processing

Psychology and cognitive neuroscience literature has long suggested predictive processing in voluntary actions filters out self-induced sensory responses to efficiently process salient sensory inputs from the periphery. This matches well with the novel peri-running neural dynamics we report, in particular the pre-running activation and suppression of VIP and Pyr neurons, respectively. Bulk manipulation of VIP/SST activity through optogenetics, as demonstrated by systems neuroscience studies^37-41,48-50^, sufficiently modulates orientation tuning and other response properties of neighbouring Pyr neurons. However, do the peri-running Pyr dynamics truly represent the visual information processing being adapted to the incoming running, or just some global effects non-specific to predictive processing? We reason that, if Pyr_Sup_ neurons are effectively recruited by predictive processing associated with running, statistically they should be biased towards the expected least informative sensory features, which, like the ‘negative image’, ought to be filtered out. In other words, the suppression of Pyr_Sup_ should be anisotropic. The post-running excitation, however, should be isotropic in order to facilitate the error reporting or the detection of novel incoming stimuli currently unknown to the brain. Our data show it is the case (Fig. 4). We managed to map orientation tuning curves of Pyr_Sup_ (n = 107) and Pyr_Post_ cells (n = 247, 8 FOVs, Fig. 4a&b) we investigated during self-initiated running (see Methods). Overall Pyr responses to drifting gratings are consistent with published results, but Pyr_Sup_ differ significantly from Pyr_Post_ cells in orientation selectivity (p = 0.012, Kolmogorov-Smirnov test, Fig. 4c). Pyr_Sup_ are more sharply tuned (OSI: 0.613 ± 0.027 vs. 0.531 ± 0.018, p = 0.012, *t*-test, Fig. 4c) and their preferred orientations (θ_pref_) are not uniformly distributed but strongly biased at 45° and 90° (p = 2.7 × 10^−15^, Pearson’s chi-squared test, Fig. 4d), supporting an anisotropic pre-running suppression. Pyr_Post_’s θ_pref_, however, follows a uniform distribution (p = 0.96, Pearson’s chi-squared test, Fig. 4d), indicating an isotropic post-running excitation. To obtain further insights, we binned θ_pref_ into 4 behaviourally cardinal pairs, which is justified by the low direction selectivity of V1 Pyr cells (Supplementary Fig. 10), and quantified θ_pref_ distributions by FOVs to control possible sampling bias. Results confirm an isotropic excitation of Pyr_Post_ neurons, as θ_pref_ of Pyr_Post_ distributes uniformly and shows no apparent preference (Fig. 4e). This is in sharp contrast with the highly anisotropic effect in Pyr_Sup_ cells: suppression readily suppress, but does not over-suppress Pyr_Sup_ tuned to 0°/180° (p = 0.99) and 135°/315° (p = 0.57, *t*-test), which is consistent with its projected function in visual flow. However, suppression is stronger to Pyr_Sup_ tuned to 90°/270° (p = 0.04), but weaker to 45°/225° (p = 0.01, Fig. 4e). Functionally this represents a re-allocation by the VIP-SST disinhibition of V1 processing power towards 45°/225°, which is behaviourally more informative to the animal because for running in the horizontal plane, an object more likely comes from top-front based on previous experience (Fig. 4f). As a result, activity tuned to 90°/270° reduces, probably due to the limited amount of computational resource in V1.

**Figure 4.**
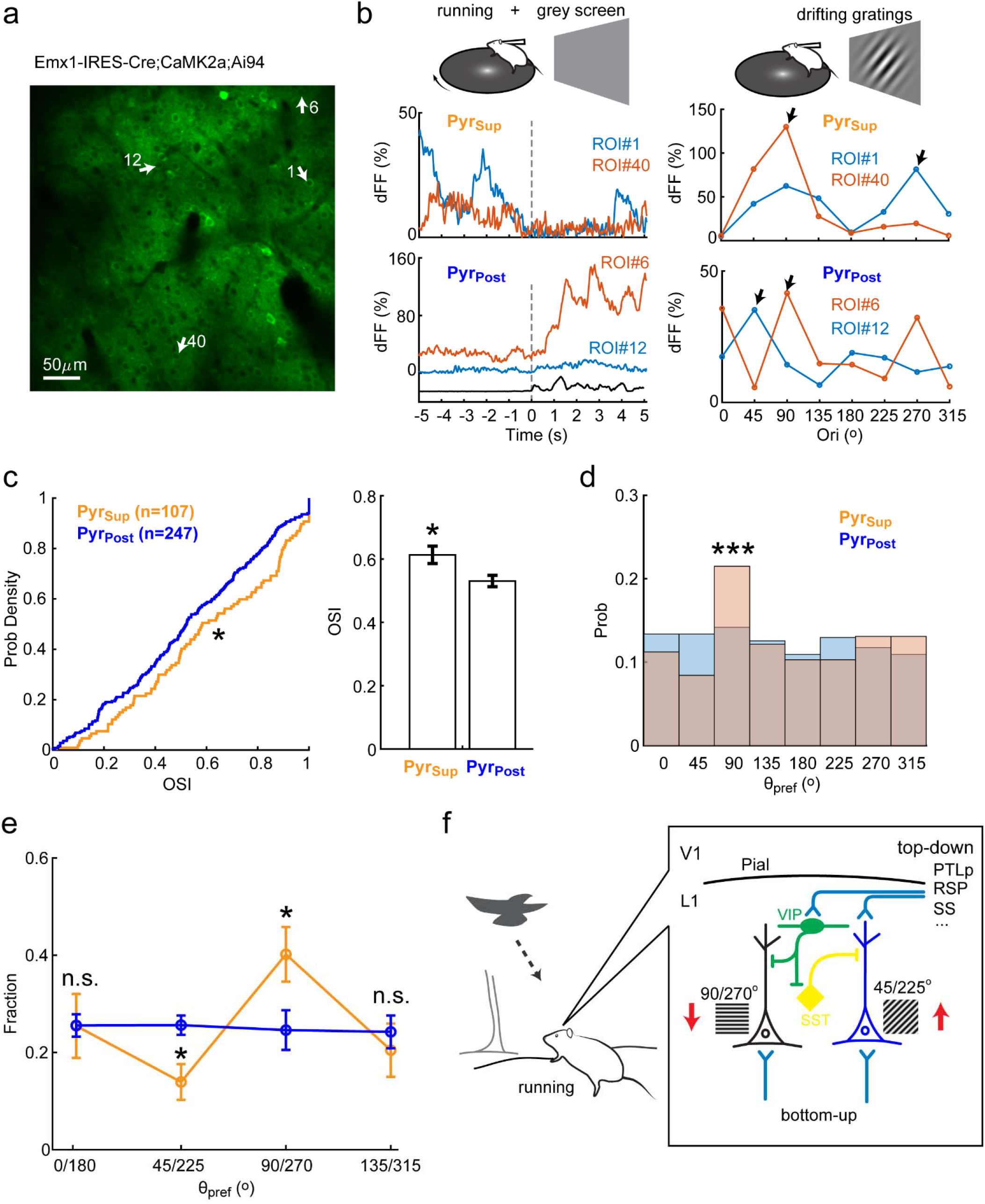
Adaptive modulation of Pyr orientation tuning by predictive processing during voluntary running. **a&b**. Orientation tuning curve mapping in the same set of Pyr neurons characterized during voluntary running. **a**. Z-projection image of the same set of Pyr cells in Fig. 2a. **b**. Orientation tuning mapping of identified Pyr_Sup_ and Pyr_Post_ cells (left column) with whole-screen drifting gratings (right column). Data are from corresponding ROIs labeled in **a**. Black arrows indicate the preferred orientation (θ_pref_). **c**. Pyr_Sup_ and Pyr_Post_ neurons significantly differ in orientation selectivity, as quantified by differences in cumulative distributions (left panel) and mean values (right panel) of OSI between Pyr_Sup_ and Pyr_Post_ cells. **d**. θ_pref_ distributions of Pyr_Sup_ and Pyr_Post_ neurons. Note θ_pref_ in Pyr_Post_ nearly follows a uniform distribution, but deviates considerably from it in Pyr_Sup_ cells, especially at 45° and 90°. **e**. Normalization of θ distributions by FOVs confirms that voluntary running anisotropically modulates Pyr_Sup_ neurons, but the post-running excitation is isotropic. **f**. Peri-running neural dynamics in the VIP-SST-Pyr microcircuit may help the brain adapt to incoming behaviorally salient sensory features (see main text for detail). *: p < 0.05, ***: p < 0.001, n.s.: not significant, Error bars: s.e.m.

To establish causal relationship, we optogenetically manipulated activities of top-down projections and VIP INs in anesthetised mouse V1 with channelrhodopsin (ChR2) and halorhodopsin, respectively^39^ (Supplementary Fig. 11, see Methods). Cell-attached recordings were conducted to compare V1 Pyr responses to drifting gratings before (ctrl) and after laser stimulations. Consistent with our Pyr_post_ data, focal activation of frontal top-down axons in V1 (top-down+) causes disinhibition of V1 Pyr responses^37-42,46^, which is completely reversed by the inactivation of VIP INs (Normalized θ_pref_ response, top-down+ vs. top-down+/VIP-, p = 0.015, Supplementary Fig. 11b). In addition, inactivating VIP INs during top-down activation, but not top-down activation alone, significantly sharpens the orientation tuning of V1 Pyr neurons (ctrl vs. top-down+/VIP-, p = 0.04, Supplementary Fig. 11c). This sharpening effect validates pre-running VIP activation suppresses sharply tuned Pyr_sup_ neurons, because blocking VIP activities during top-down activation would release those Pyr neurons from suppression thus result in a sharpening effect of Pyr tuning curves, which is exactly what the optogenetic data show. Functionally Pyr_sup_ suppression may enhance behavioural performance^51^. Due to the randomness of self-initiated running, as we have pointed out, it remains infeasible to present optogenetic manipulation before running to examine the running-dependent anisotropic suppression (Fig. 4f). In summary, these results experimentally verify the causal role of the VIP-SST-Pyr microcircuit in predictive processing and provide pivot *in vivo* evidence for neurobiological substrate of predictive processing.

## Discussion

In the present study, we have dissected the neuronal cell-type and circuit basis for predictive processing, which is a fundamental concept in cognitive neuroscience, psychology, cybernetics and artificial intelligence yet with limited systems neuroscience knowledge^1-11,52^. Employing a self-initiated running paradigm, we discover robust VIP activation prior to running thus reveal a novel neuronal signature of visual predictive processing in voluntary action. Preparatory activation in V1 is selective to VIP INs, suggesting the pivot role of VIP INs in top-down and bottom-up crosstalk of predictive processing. We further identify that, with converging anatomical and functional evidence, the VIP-SST-Pyr microcircuit is the key element of predictive processing due to the fact that it is the recipient of long-range top-down projections from the ‘upper layer’ brain regions known to fire preparatory neural activities and its unique peri-running dynamics preadapt V1 Pyr neurons to the incoming running, which matches well with theoretical projections. Together, these findings illustrate the neural substrate of predictive processing, thus bridge the long-existing gap between psychology/cognitive neuroscience and systems neuroscience.

The SST-VIP circuit has been previously suggested for visual mismatch signal (prediction-error) processing^47^, yet the upstream neural circuitry of predictive processing remains unknown. Our data of the VIP-SST-Pyr circuit complete this missing link and complement the SST-centred circuitry for prediction-error plasticity during development. VIP-SST motif plays critical roles in sensory information processing (e.g. gain control), selective attention and associative learning^37-41,48-50^. Here we present novel evidence for its function in voluntary actions. VIP-SST-Pyr microcircuit may universally underlie neural computations for predictive processing, our results thus will help to generate better understandings of the neurobiology of cognitive functions important to human nature such as sense of agency/volition^53^. Since the seminal work of the ‘Readiness Potential’ in human supplementary motor area, preparatory neural activity has been found relevant to voluntary action planning, decision-making and time sensation^2-20^. The identification of the VIP-SST-Pyr mirocircuit could shed new light on pinpointing neural substrate of sophisticated brain functions for more fine-grained, mechanistic knowledge. Future studies are required to elucidate genetic, molecular and connective signatures of VIP/SST subgroups to allow better-targeted manipulations of higher-order cognitive functions, in order to develop novel treatments on psychiatric disorders without undesired confounding effects^12^.

## Acknowledgements

This work was supported by grants from National Natural Science Foundation of China NSFC 31871055 (L.L.), NSFC 31871051 (S.Y.Z), Guangdong Science and Technology Department 2017B030314026 (L.L.) and 2018B030334001 (L.L.). We thank Dr. Karel Svoboda and his laboratory for technical assistance. The authors wish to thank the Allen Institute founders, Paul G. Allen and Jody Allen, for their vision, encouragement and support.

## Author Contributions

L.L. conceived and supervised the project. L.H. and L.L. performed Ca imaging experiments. L.L. and J.Z. analyzed Ca imaging data. L.Z.W. and S.Y.Z. conducted the retrograde rabies tracing and optogenetics experiments. Y.Y.X., C.C.L. and Q.L.P. helped Ca imaging data analysis. L.L wrote the paper with inputs from all authors.

## Data availability

The authors declare that all data supporting the findings of this investigation are available within the article, its Supplementary figures, and from the corresponding authors upon reasonable request.

## Methods

Experimental procedures were in accordance with NIH guidelines and approved by the Institutional Animal Care and Use Committees (IACUC) of Sun Yat-sen University, Shanghai Jiao Tong University School of Medicine and Allen Institute for Brain Science. In the current study, 2-p Ca imaging were performed in adult transgenic mice (2 – 7 months, both sex), including VIP-IRES-Cre;Ai162 (GCaMP6s, n = 6), Emx1-IRES-Cre;CaMK2a-tTA;Ai94 (GCaMP6s, n = 3) and SST-IRES-Cre;Ai163 (GCaMP6s, n = 5). Monosynaptic retrograde tracing was conducted with VIP-IRES-Cre mice (n = 4). Optogenetic manipulation was done with VIP-IRES-Cre;Ai39 mice (Halo, n = 9).

### Viral injections

Viral vectors were prepared as published previously^36,39^. For retrograde monosynaptic tracing Glycoprotein-deleted (ΔG) and EnvA-pseudotyped rabies virus (RV-ΔG-EGFP+EnvA, 2.4 × 10^8^ IU/mL), AAV2-CAG-DIO-TVA-mCherry (2.1 × 10^13^ gc/mL) and AAV2-CAG-DIO-Glycoprotein (8.7 × 10^12^ gc/mL) were used. The genomic titer of AAVs and RV was determined by quantitative polymerase chain reactions. Both AAVs and RV were produced by BrainVTA. In order to trace presynaptic inputs of VIP neurons in V1, AAV2-CAG-DIO-TVA-mCherry and AAV2-CAG-DIO-Glycoprotein (1:2, 500 nL) were co-injected into V1 (3 mm posterior and 2.5 mm lateral to Bregma, depth 0.5 mm) of VIP-Cre mice. This resulted in cell-type specific expressions of TVA receptor and rabies glycoprotein, which are required for virus infection and trans-synaptic spread, respectively, in Cre-positive VIP neurons. Two to three weeks later, RV-ΔG-EGFP+EnvA (500 nL) was injected into the same site as AAV injection. Histology was conducted 7 days after rabies virus injection. For optogenetics, AAV2-CaMKII-hChR2(H134R)-EYFP (UNC Vector Core) and AAV-DJ-CaMKII-hChR2(H134R)-EYFP (Stanford Neuroscience Gene Vector and Virus Core) were used. To activate top-down projections in V1, AAV2-hChR2(H134R) or AAV-DJ-hChR2(H134R) was injected into ipsilateral frontal cortex (800 nL, 0.2 mm anterior and 0.3 mm lateral to Bregma, depth 0.9 mm).

### Surgery

Detailed surgical procedures have been published elsewhere^35^. Briefly, a custom-made, pre-notched metal head-post was implanted under Isoflurane anesthesia (1.5 -2% in O_2_). A circular craniotomy was performed over left visual cortical area followed by durotomy. Silastic sealant (Kwik Sil) was applied to the craniotomy before a 3 mm X 3 mm polycarbonate coverslip was placed over the exposed cortical area centering on 1.3 mm anterior and 3.1 mm lateral to the Lambda. Notches on the head-post were carefully aligned during surgery to establish a coordinate system for V1 positioning in Ca imaging. The headpost and coverslip were cemented firmly onto the skull. After 7 days of recovery, mice were habituated to head fixation and presentation of visual stimulations on an in-house made running device (a freely-rotating running disc) 2 hours per day for 1 week before imaging experiments started. During the daily habituation, mice were free to rest or run on the running disc, but not specifically trained for continuous locomotion.

### *In vivo* 2-p Ca Imaging

Ca imaging was performed using a Bruker 2-p microscope equipped with 8 kHz resonant scanners and a Chameleon Ultra II Ti:Sapphire femtosecond laser system (Coherent). The microscope was within a light-proof Faraday cage, which sufficiently blocked ambient light from surrounding laboratory area when the front doors were shut. Fluorescence was excited at 920 nm, collected into two spectral channels using green (510/42 nm) and red (641/75 nm) emission filters and saved on a local computer. Imaging was conducted ∼150 – 300 µm underneath the pial surface with a 16× water-immersion objective lens (Nikon, NA 0.8). Image data were acquired at 512 × 512 pixels, 30 Hz frame rate with laser power <70 mW measured after objective to ensure image quality. During imaging, mice were awake, head-fixed and allowed to run or rest on the running disc ‘at will’ (Fig. S1b), with or without visual stimulations. For imaging in darkness, in addition to turning off the visual stimulation monitor and covering all light emission spots within the Faraday cage, a custom-made black light shield, which fitted tightly with the headpost, was installed before imaging to fully cover the objective.

In a typical imaging experiment, we first imaged Ca activities within the 400 × 400 µm field of view (FOV) independent of running occurrence for 200 sec without presentation of structured visual stimuli. A grey screen was presented during imaging on a calibrated LCD screen at 50% contrast to the contralateral eye, unless imaging was conducted in darkness. Orientation tuning mapping might follow after a ∼5 – 10 minute break and was conducted to the same FOV with drifting grating stimuli (see below). Vasculature landmarks and coordinates calculated from headpost notches were used to keep FOVs from overlapping. To ensure animal’s health, total imaging time in the same animal on the same day was less than 2 hours.

### *In vivo* cell-attached recording

Mice were anesthetized with urethane (0.6-1.2 g/kg, IP) and chlorprothixene (5 mg/kg, IM) after at least 2 weeks of ChR2 expression. During experiment, body temperature of the animal was maintained at 37 °C using a feedback-controlled heating pad. A craniotomy was made over V1 after firmly cementing a custom head-post onto the skull. V1 Pyr neurons were recorded with borosilicate micropipettes (∼4-8 MΩ) filled with HEPES buffered ACSF under cell-attached configuration. Signals were acquired with a Multiclamp 700B amplifier, filtered between 0.5-2 kHz, sampled at 10 kHz and stored on a local PC using the Clampex software. ACSF was frequently applied to the craniotomy to prevent the exposed cortex from drying.

### Visual stimulations

Whole-screen sinusoidal drifting gratings were generated using Psychopy software and presented on the calibrated LCD monitor. The centre of the monitor was positioned ∼22 cm away from the centre of the contralateral eye. The stimulus set consisted of 8 orientations (0° - 315°, 45° increment), 4 spatial frequency (SF, [0.02, 0.04, 0.08, 0.16] cycle per degree, cpd) and 1 temporal frequency (2 Hz). Gratings were presented to the animal at 80% contrast for 2 seconds, separated by 2 seconds of grey screen at mean illuminance and repeated 5 times in a random sequence. For optogenetics experiments, drifting gratings were presented for 4 sec. A grey screen at mean illuminance was randomly inserted and presented for 15 times. Due to the choice of whole-screen visual stimuli, we didn’t map the receptive field or tract the pupil position.

### Optogenetics

ChR2-expressing axons of frontal top-down projections were focally activated using blue laser pulses (wavelength 473 nm, 10 Hz, 5 ms) delivered via an optic fiber (200 µm in diameter) over V1. The power of laser was measured at ∼3-5 mW at the open end of the optic fiber. To inactivate Halo-expressing V1 VIP INs, an optic-fiber (600 µm in diameter) was used to deliver yellow laser (wavelength 593 nm, 3 sec, power of 5-8 mW) over V1. Pyr responses to whole-screen drifting gratings were compared between no laser (ctrl), blue laser (top-down+) and blue+yellow lasers (top-down+/VIP-) conditions, representing control, top-down activation (resembling running) and inactivation of V1 VIP INs during running, respectively. The open ends of optic fibers were held in place over the recorded Pyr neurons and in close proximity to the exposed V1 surface by micromanipulators.

### Histology

Detail of histology has been published elsewhere^36^. After embedding and freezing, the brain was sectioned into 50-μm coronal slices using a cryostat. One out of every three sections were imaged in the high-throughput slide scanners (Nanozoomer-2.0RS) for further processing. A custom written software package was used to analyze the digitized brain images. The analysis software consists of four modules: atlas rotation, image registration, signal detection, and quantification/visualization. The Allen Mouse Brain Atlas was rotated to mimic the aberrant sectioning angle of the experimental brain. Images of brain sections were aligned to the rotated reference atlas for further quantification. The detection module was used to record the position of manually identified EGFP-labeled neurons in each digitized brain section image. After detection and registration, signals were quantified across the whole brain. Since the number of labeled neurons varied across brains, the input density from each region was quantified by dividing the number of labeled neurons found in that region by the total number of labeled neurons detected in the entire brain, with the exception of the injection site.

### Data analysis

Behaviours of head-fixed mice during Ca imaging were quantified to validate the behavioural paradigm employed in the current study was adequate to study neural correlates of self-initiated (voluntary) actions. Data of running velocity were analyzed and results were summarized in Supplementary Figs. S1b, S7b and S9b, respectively. Firstly, in our experiments running was sporadic, i.e. mice didn’t necessarily run in every imaging session. Secondly running was typically brief, only occurred in ∼5% - 10% of the total imaging time. This distinguished the current study from those that specifically trained animals to run continuously, which engaged the neuromodulatory system profoundly. Thirdly, the start of running was random in time, which meant animals could initiate running at any time unknown to the experiment, in a way similar to the ‘at will’ performance in human studies. This was because running onset followed a uniform distribution, which made it impossible to deliver temporally coupled manipulation, and at the same time, directly argued against the possibility that the running represented a learned behaviour in response to head-fixation or was driven by external sensory cue. Finally, in our dataset, running exhibited these features independent of transgenic mouse lines or imaging conditions. We want to emphasize that our experiment design, especially the use of naïve mice in absence of external sensory stimuli, ensures minimal engagement of selective attention and associative learning processes. Therefore, we concluded that the running behaviour in our paradigm was self-initiated and peri-running neuronal dynamics accurately reflected neural processes underlying voluntary running.

Ca imaging data were analyzed using in-house Matlab scripts. Green image stacks were first corrected for in-plane motion artifacts using published cross-correlation motion correction method between frames^54^. Image segmentation was done with custom-made Matlab package^34^. Regular ring-shape ROIs were defined and neuropil subtraction was conducted for Pyr cells imaged from Emx1-IRES-Cre;CaMK2a-tTA;Ai94 mice as described previously^34^. For inhibitory VIP and SST interneurons, as we reported before, GCaMP6s expression in VIP-IRES-Cre;Ai162 and SST-IRES-Cre;Ai163 mice showed no clear nucleus exclusion^33^, patch ROIs were thus used for VIP and SST interneurons and green fluorescence signals were computed from all pixels within the patch ROI^34^. Due to the sparseness of cortical inhibitory interneurons, neuropil signal was not corrected for VIP and SST neurons^34^.

For imaging data of voluntary running, ΔF/F was calculated for each ROI with the mode of raw F traces F_mod_, i.e. ΔF/F = (F - F_mod_)/F_mod_. Fluorescently active ROIs were then determined by threshoulding the ΔF/F traces at 3-4 standard deviations above the mean with manual verifications. Inactive ROIs were rejected from further analysis. To analyze peri-running neuronal dynamics, running velocity were threshoulded at 1 cm/s (ref #44). The time points at which the velocity up- and down-crossed the 1 cm/s threshould defined running onsets and offsets, respectively. Consistent with previous results^42,44^, 1cm/s is at the low end of the velocity distribution. For better temporal resolution, velocity data were not smoothed. Imaging sessions that failed to reach the velocity threshould were considered no running and excluded from further analysis. Next for each imaging session, we identified running episodes during which the mice remained stationary (i.e. no suprathreshould events detected) for at least 10 sec before the running onset, in order to control the slow decay of GCaMP6s. ΔF/F traces of active ROIs within -10s to 5s from the running onset were extracted (using the imaging frame nearest to the running onset as time 0) and averaged across running episodes, in order to control the variabilities between running episodes. The averaged ΔF/F traces were carefully examined with the running data for suspicious pre-running subthreshould movement. Data showing ΔF/F deflection corresponding with the subthreshould movement were excluded from analysis, unless could be proved otherwise (e.g. Fig. 1c&d). For individual ROI, latency was determined from running-episode averaged ΔF/F trace as the earliest time point (in the [-3s, 2s] range) above the mean plus 3 - 4 times standard deviation, both computed from the [-10s, 0s] baseline data. In a small number of cells (<5%), baseline was slightly adjusted, e.g. to [-10s, -1s] or [-10s, -2s], to deal with the excessive firing just before the running initiation. Cells with negative and positive latencies were classified as ‘Pre’ and ‘Post’ cells, respectively. Those cells that failed to meet the latency criteria but had mean ΔF/F within [0s, 5s] lower than mean baseline ΔF/F were categorized into the suppressed-by-running group (‘Sup’). The rest was considered non-responsive cells. We found this resulted in complete, non-overlapping classification of VIP, SST, Pyr neurons. To control the variation of post-running activities among individual neurons, ΔF/F was normalized by the group average of ΔF/F during corresponding baseline activities. We also normalized ΔF/F individually by the average of baseline activity, which controlled baseline variations, and results were consistent (data not shown). Due to the slow decay of GCaMP6s, we didn’t analyze ΔF/F activities for running offsets.

To analyze visually evoked Ca responses, ΔF/F was calculated for individual trials. The average F signal within 2 sec immediately before the stimulus onset were used as the baseline. In case that companion imaging sessions of voluntary running were available, ROIs were imported from the corresponding running session, adjusted for any spatial shifts to ensure best alignment. For visual response analysis, fluorescently active ROIs were analyzed independent of the occurrence of running. ΔF/F was averaged over the 2 sec stimulus duration, then by stimulus conditions. To calculate the orientation selectivity index (OSI), the preferred orientation (θ_pref_) was determined by the orientation that evoked the strongest response at the preferred SF. OSI was calculated as OSI = (R_pref_ - R_orth_)/(R_pref_ + R_orth_), where R_pref_ and R_orth_ stands for the response magnitude at θ_pref_ and the orthogonal orientation (θ_pref_ + π/2), respectively. Direction selectivity index (DSI) was calculated as DSI = (R_pref_ - R_oppo_)/(R_pref_ + R_oppo_), where R_oppo_ stands for the response magnitude at the opposite direction (θ_pref_ + π).

Cell-attached data were analyzed using in-house scripts. Only recordings stable for more than 5 min and with sufficiently high seal resistance (10-100 MΩ) were included in final data analysis.

It has been reported that some Emx1-IRES-Cre;CaMK2a-tTA;Ai94 mice could have aberrant epileptiform activity, which may confound our Pyr data. No signs of seizure-like behavior were noticed either during imaging or in home-cages from the Ai94 mice included in our data analysis. Running statistics of these Ai94 mice showed no difference from those VIP and SST transgenic mice we imaged, and Pyr responsiveness was fully comparable with that previously reported in normal mice. All these pieces of evidence suggested no behavioral abnormality in these three Ai94 mice, thus our Ai94 results were NOT confounded by epileptiform activity that may occur in some Ai94 mice.

