## supplementary information for "Neuronal cell-type and circuit basis for visual predictive processing"

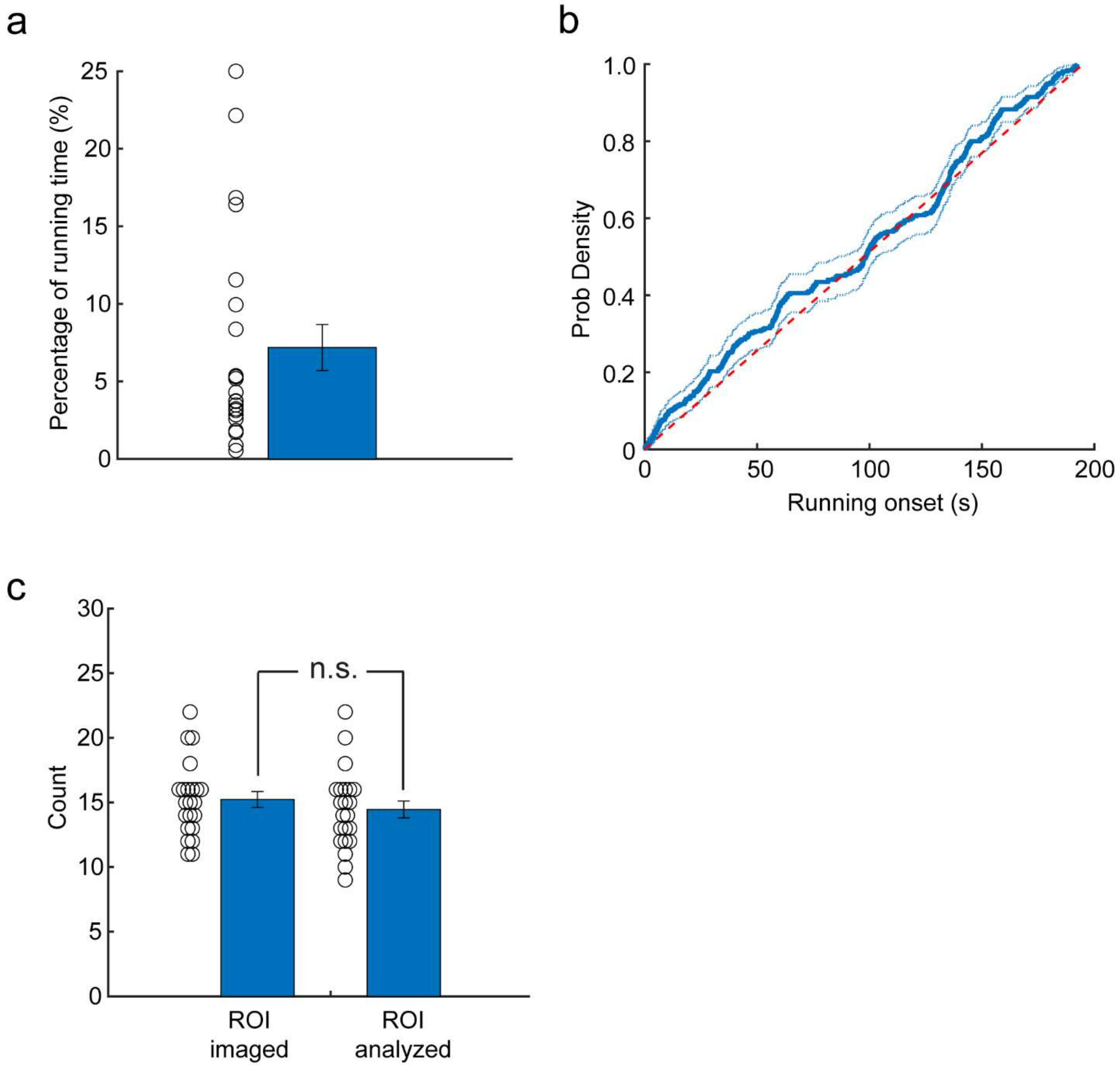


**Supplementary Figure 1. Brief, sporadic voluntary running in awake, head-fixed VIP-IRES-Cre;Ai162 mice. a**. Percentage of running time (7.2 ± 1.5%, mean ± s.e.m., same below) in 22 FOVs analyzed. **b**. Cumulative distribution of the running onset along the 0 - 195 sec imaging time. Probability density (thick line) and confidence interval (dotted line) are both plotted. Red dashed line stands for the uniform distribution. Nearly random occurrence of running events indicates the running behaviour is self-initiated, neither an evoked response to external sensory stimuli nor a learned behavior associated with the imaging setup. **c.** No ROI selection bias in data analysis. No significant difference between numbers of total ROI imaged (15.2 ± 0.6, left) and fluorescently active ROI analyzed (14.5 ± 0.7, right, p = 0.40, *t*-test) per FOV in VIP-Cre;Ai162 mice. On average 95.1 ± 2.2% VIP neurons are fluorescently active and included in our final analysis. n.s.: not significant, Error bars: s.e.m.

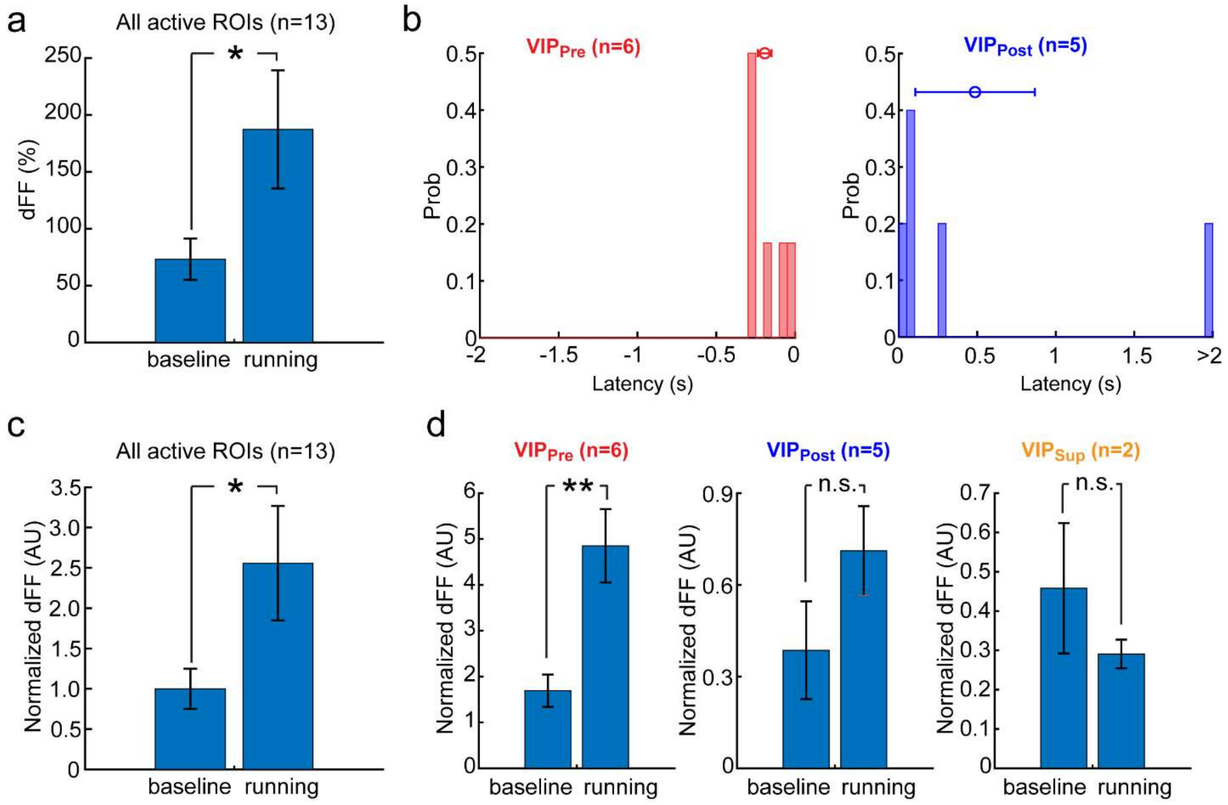


**Supplementary Figure 2. Extended data of peri-running dynamics of example VIP neurons in Fig. 1. a**. Significant increase of mean ΔF/F activity of example VIP neurons in Fig. 1b to f during running (baseline vs. running: 73.2 ± 18.2 vs. 187.3 ± 52.0, n = 13, p = 0.049, *t*-test). **b.** Latency histograms of example $VIP_{Pre}$ (left) and $VIP_{Post}$ neurons (right). See main text for individual mean and s.e.m. values. **c&d.** Normalized ΔF/F activities during baseline imaging and running in all example VIP (**c**), $VIP_{Pre}$ $VIP_{Post}$, and $VIP_{Sup}$ neurons (**d**). Running significantly increases overall VIP activity (1.00 ± 0.25 vs. 2.56 ± 0.71, n = 13, p = 0.049). During running ΔF/F also significantly increases in $VIP_{Pre}$ (1.69 ± 0.35 vs. 4.85 ± 0.80, n = 6, p = $4.7 \times 10^{-3}$), but has insignificant effects in $VIP_{Post}$ (0.39 ± 0.16 vs. 0.71 ± 0.15, n = 5, p = 0.17) and $VIP_{Sup}$ (0.46 ± 0.17 vs. 0.29 ± 0.04, n = 2, p = 0.43) neurons. * p<0.05, ** p<0.01, n.s.: not significant, Error bars: s.e.m.

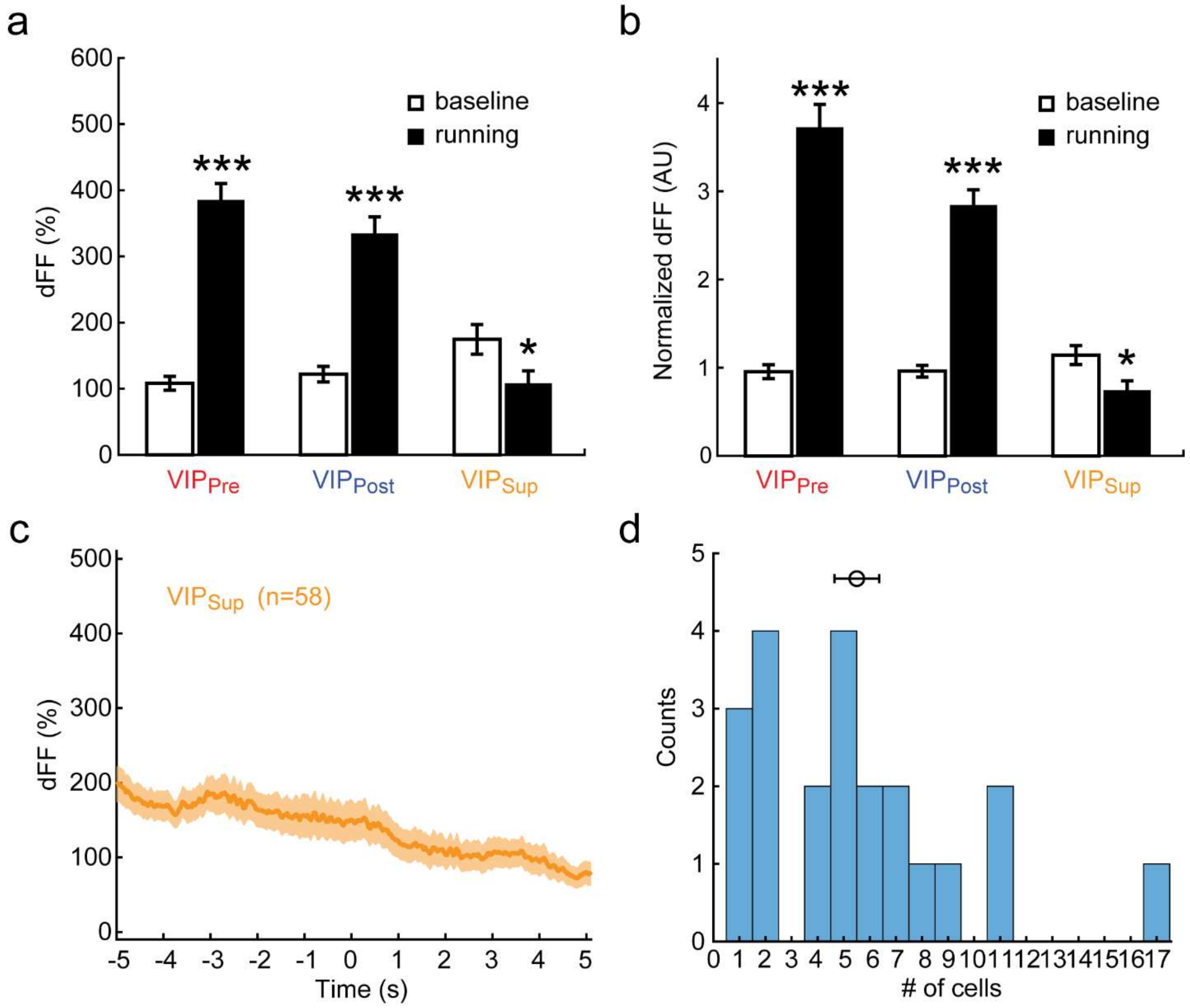


**Supplementary Figure 3. Population analysis of peri-running dynamics of VIP neurons. a**. Mean ΔF/F activities during baseline imaging and running in $VIP_{Pre}$, $VIP_{Post}$ and $VIP_{Sup}$ neurons. Both $VIP_{Pre}$ (baseline vs. running: 108.1 ± 10.5 vs. 384.5 ± 25.6, n = 121, p = 7.0 × $10^{-20}$, *t*-test) and $VIP_{Post}$ (122.0 ± 11.6 vs. 334.0 ± 25.6, n = 139, p = 7.3 × $10^{-13}$) neurons significantly increase activities after running initiation. $VIP_{Sup}$ cells, however, show significant activity suppression (174.5 ± 22.5 vs. 107.4 ± 19.7, n = 58, p = 0.027) by running. **b.** Normalized ΔF/F activities during baseline and running in $VIP_{Pre}$, $VIP_{Post}$ and $VIP_{Sup}$ neurons. Running significantly increases ΔF/F activities in both $VIP_{Pre}$ (0.95 ± 0.08 vs. 3.72 ± 0.26, p = 5.5 × $10^{-20}$) and $VIP_{Post}$ neurons (0.96 ± 0.07 vs. 2.84 ± 0.18, p = 1.2 × $10^{-19}$), but suppresses $VIP_{Sup}$ cells (1.14 ± 0.11 vs. 0.74 ± 0.11, p = 0.01). **c.** Time course of peri-running dynamics of $VIP_{Sup}$ cells. **d.** Histogram of # of $VIP_{Pre}$ INs identified per FOV (n = 22). * p<0.05, *** p<0.001, Error bars: s.e.m.

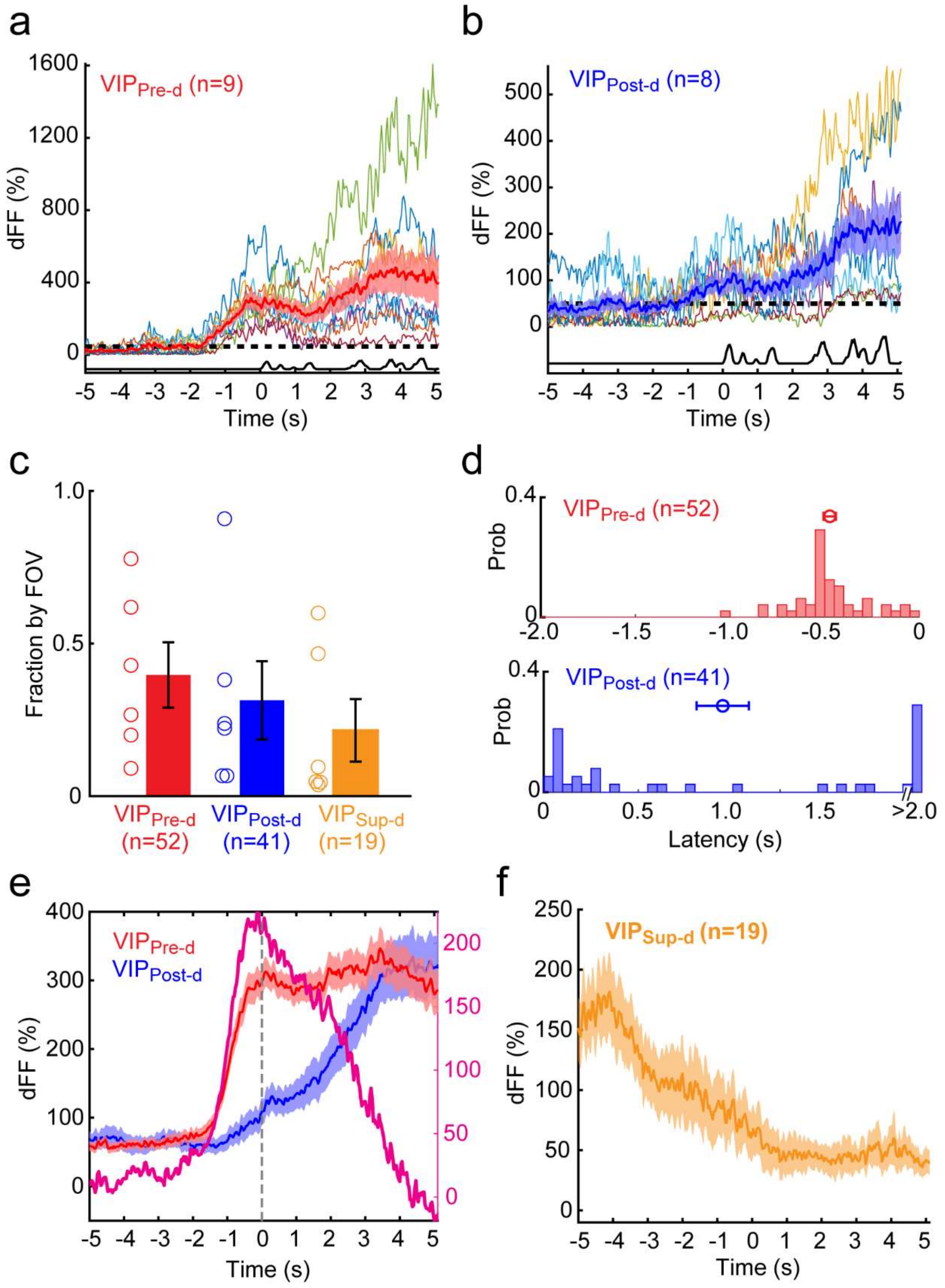

a
VIP Pre-d (n=9)
dFF (%)
Time (s)
b
VIP Post-d (n=8)
dFF (%)
Time (s)
c
Fraction by FOV
VIP Pre-d (n=52)
VIP Post-d (n=41)
VIP Sup-d (n=19)
d
VIP Pre-d (n=52)
Prob
VIP Post-d (n=41)
Prob
Latency (s)
>2.0
e
VIP Pre-d
VIP Post-d
dFF (%)
Time (s)
f
VIP Sup-d (n=19)
dFF (%)
Time (s)

**Supplementary Figure 4. Robust pre-running activation of V1 VIP INs imaged in darkness. a&b.** Time courses of peri-running activities of $VIP_{Pre\text{-}d}$ (n = 9, **a**) and $VIP_{Post\text{-}d}$ neurons (n = 8, **b**) in an example FOV of total 21 ROIs imaged in darkness. The rest are one $VIP_{Sup\text{-}d}$ and 3 non-responsive VIP INs. Note $VIP_{Post\text{-}d}$ activity also increases before running initiation, but not significantly. **c – e.** Population data of VIP INs (112 active ROIs, 6 FOVs in3 VIP-Cre;Ai162 mice) imaged in darkness showing fractions of $VIP_{Pre\text{-}d}$ (39.7 ± 10.7%, total 52/112), $VIP_{Post\text{-}d}$ (31.4 ± 12.9%, total 41/112) and $VIP_{Sup\text{-}d}$ (20.2 ± 10.7%, total 19/112) neurons by FOV (**c**); latency distributions of $VIP_{Pre\text{-}d}$ (-0.473 ± 0.029 sec) and $VIP_{Post\text{-}d}$ (0.947 ± 0.138 sec) neurons (**d**); average time courses of $VIP_{Pre\text{-}d}$, $VIP_{Post\text{-}d}$ populations and net pre-running activation (**e**). Consistent pre-running activation of VIP INs are found before the mice start running in darkness, ruling out confounding influences by uncontrolled visual input. **f.** Time course of peri-running dynamics of $VIP_{Sup\text{-}d}$ cells.

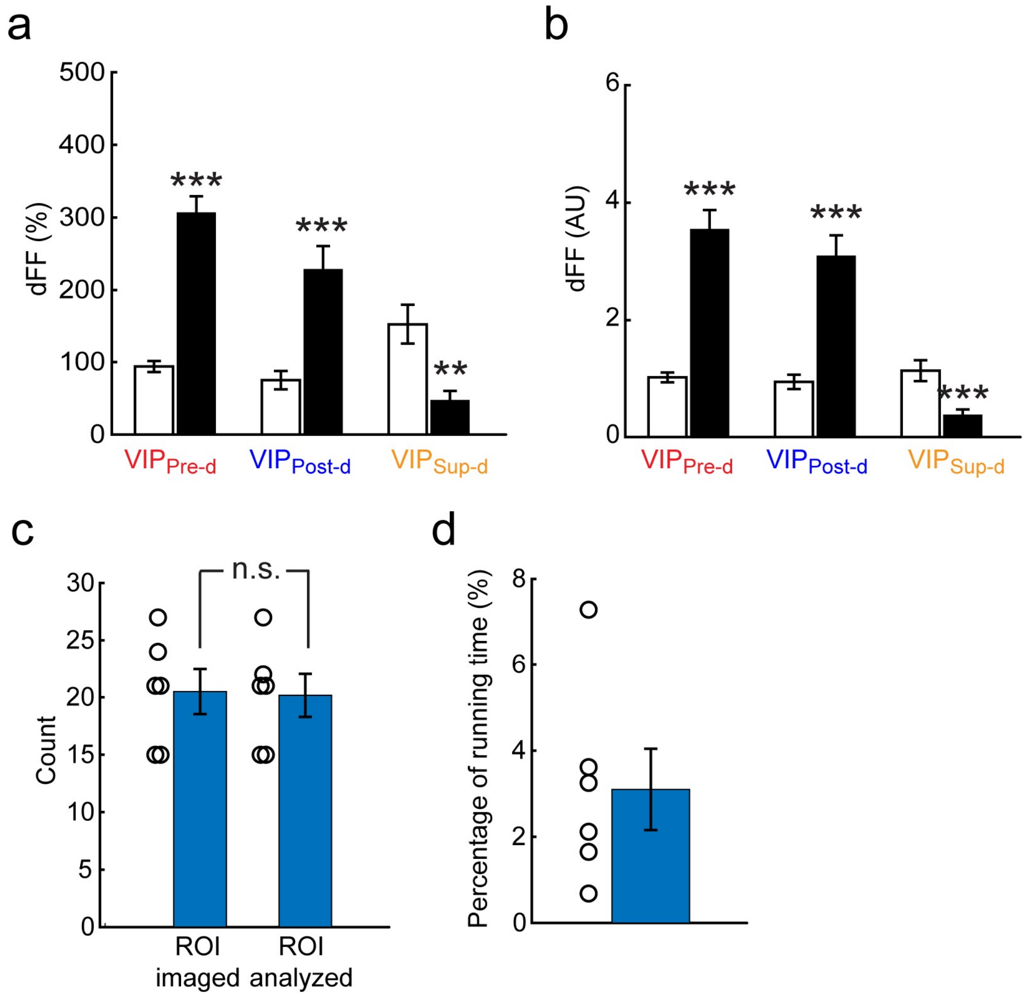
a
dFF (%)
500
400
300
200
100
0
***
***
**
VIPPre-d
VIPPost-d
VIPSup-d
b
dFF (AU)
6
4
2
0
***
***
***
VIPPre-d
VIPPost-d
VIPSup-d
c
Count
30
25
20
15
10
5
0
n.s.
ROI imaged
ROI analyzed
d
Percentage of running time (%)
8
6
4
2
0

**Supplementary Figure 5. Extended analysis of V1 VIP INs imaged in darkness. a**. Mean ΔF/F activities during baseline imaging and running in $VIP_{Pre-d}$, $VIP_{Post-d}$ and $VIP_{Sup-d}$ neurons. Both $VIP_{Pre-d}$ (baseline vs. running: 93.9 ± 7.6 vs. 306.3 ± 22.7, n = 52, p = $2.6 \times 10^{-14}$, *t*-test) and $VIP_{Post-d}$ (75.1 ± 12.7 vs. 228.2 ± 32.4, n = 41, p = $3.3 \times 10^{-5}$) neurons significantly increase activities after running initiation. $VIP_{Sup-d}$ cells, however, show significant activity suppression (152.7 ± 27.1 vs. 47.1 ± 13.2, n = 19, p = 0.001) by running. **b.** Normalized ΔF/F activities during baseline and running in $VIP_{Pre-d}$, $VIP_{Post-d}$ and $VIP_{Sup-d}$ neurons. Running significantly increases ΔF/F activities in both $VIP_{Pre-d}$ (1.02 ± 0.08 vs. 3.55 ± 0.33, p = $3.6 \times 10^{-11}$) and $VIP_{Post-d}$ neurons (0.94 ± 0.12 vs. 3.09 ± 0.36, p = $1.8 \times 10^{-7}$), but suppresses $VIP_{Sup-d}$ cells (1.13 ± 0.18 vs. 0.38 ± 0.10, p = $6.4 \times 10^{-4}$). **c.** No selection bias in peri-running analysis of VIP INs imaged in darkness. Numbers of ROI imaged (20.5 ± 2.0) per FOV are not significantly different from those of fluorescently active ROI analysed (20.2 ± 1.9, p = 0.90, *t*-test) in VIP-IRES-Cre;Ai162 mice. On average 98.6 ± 0.1% of imaged VIP neurons are included in data analysis. **d**. Percentage of running time (3.1 ± 0.9%) in 6 FOVs analyzed. * p<0.05, ** p<0.01, *** p<0.001, Error bars: s.e.m.

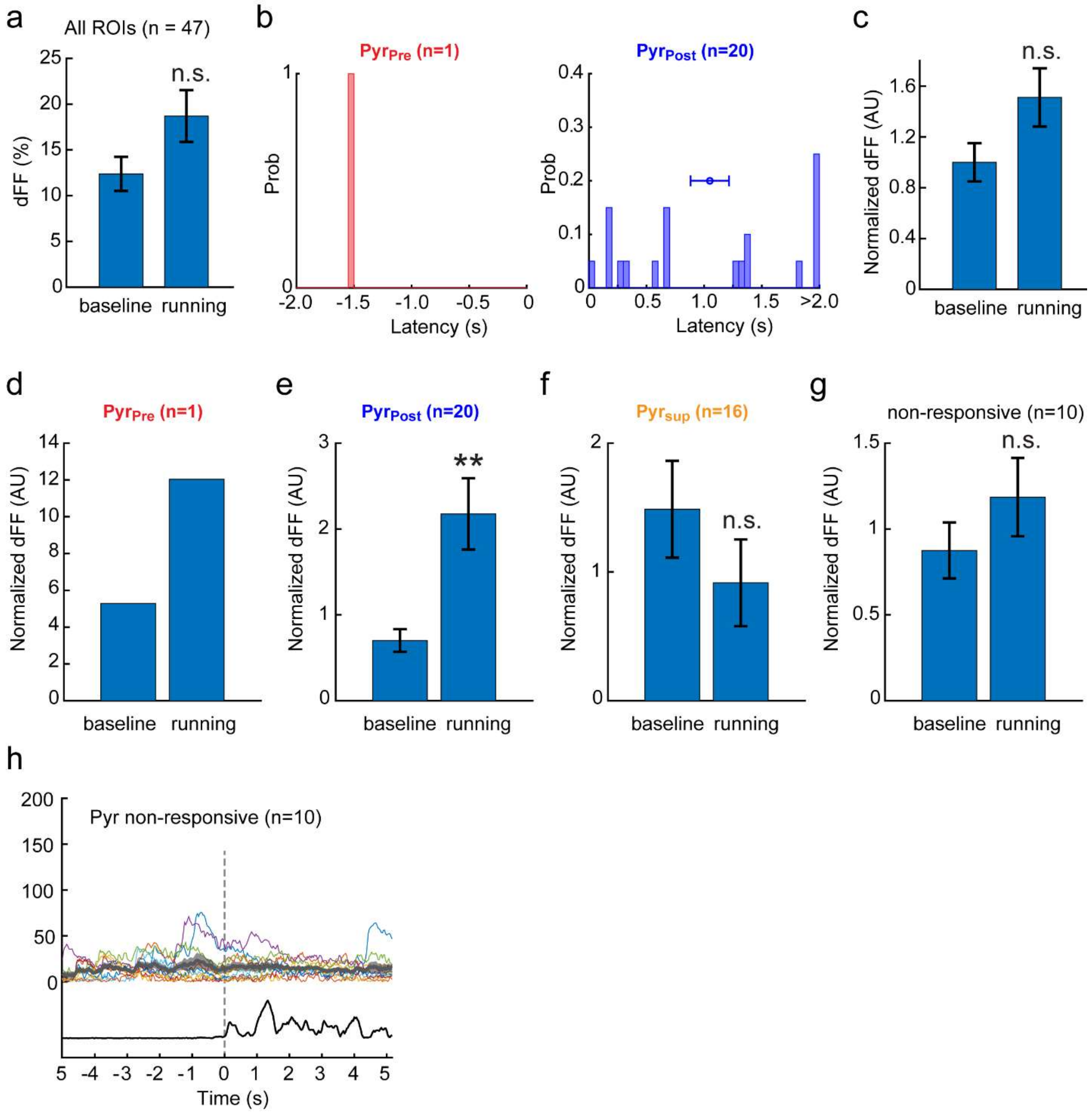


**Supplementary Figure 6. Extended data of example Pyr neurons in Fig. 2. a**. Mean ΔF/F activities of 47 example Pyr neurons shown in Fig. 2a-d. Compared with the pre-running baseline (12.4 ± 1.9), ΔF/F increases during running, but insignificantly (18.7 ± 2.8, n = 47, p = 0.07, *t*-test). **b.** Latency histograms of example $Pyr_{Pre}$ (-1.52 sec, n = 1, left) and $Pyr_{Post}$ neurons (1.049 ± 0.167 sec, n = 20, right). **c - g.** Normalized ΔF/F activities between baseline and running in all example Pyr (**c**), $Pyr_{Pre}$ (**d**), $Pyr_{Post}$ (**e**), $Pyr_{Post}$ (**f**) and non-responsive Pyr neurons (**g**). Running significantly increases $Pyr_{Post}$ activities (baseline vs. running: 0.70 ± 0.13 vs. 2.18 ± 0.42, p = 0.002, *t*-test). **h.** Peri-running dynamics of non-responsive Pyr neurons. ** p<0.01, n.s.: not significant, Error bars: s.e.m.

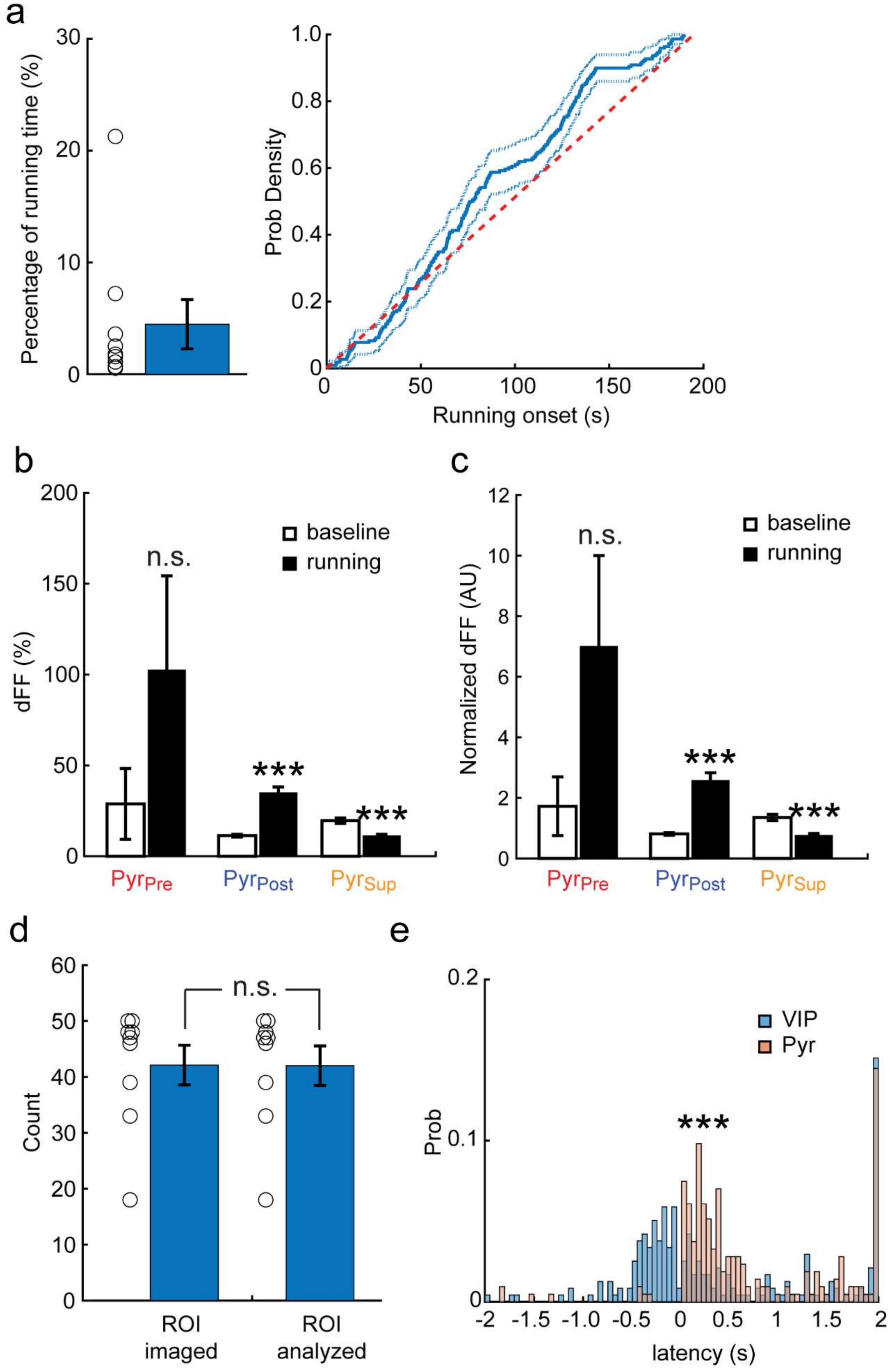
a
Percentage of running time (%)
0
10
20
30
Prob Density
0
0.2
0.4
0.6
0.8
1.0
0
50
100
150
200
Running onset (s)
b
dFF (%)
0
50
100
150
200
n.s.
***
***
baseline
running
PyrPre
PyrPost
PyrSup
c
Normalized dFF (AU)
0
2
4
6
8
10
12
n.s.
***
***
baseline
running
PyrPre
PyrPost
PyrSup
d
Count
0
10
20
30
40
50
60
n.s.
ROI
imaged
ROI
analyzed
e
Prob
0
0.1
0.2
VIP
Pyr
***
-2
-1.5
-1.0
-0.5
0
0.5
1
1.5
2
latency (s)

**Supplementary Figure 7. Population analysis of peri-running dynamics in Pyr neurons. a**. Running statistics in Emx1-IRES-Cre;CaMK2a-tTA;Ai94 mice. Left: percentage of running time (4.5 ± 2.2%) in 9 FOVs analyzed. Right: cumulative distribution of the running onset. **b**. Mean ΔF/F activities during baseline imaging and running in $Pyr_{Pre}$, $Pyr_{Post}$ and $Pyr_{Sup}$ neurons. $Pyr_{Post}$ neurons significantly increase activities after running starts (baseline vs. running: 11.3 ± 0.8 vs. 34.8 ± 3.3, n = 252, $p = 1.2 \times 10^{-11}$, *t*-test). $Pyr_{Sup}$ cells, however, show significant activity suppression (19.6 ± 1.5 vs. 11.2 ± 0.9, n = 118, $p = 2.9 \times 10^{-6}$). $Pyr_{Pre}$ only show insignificant change (28.8 ± 19.5 vs. 102.4 ± 51.9, n = 8, p = 0.21), due to its limited sample size. **c**. Normalized ΔF/F activities during baseline and running in $VIP_{Pre}$ (1.72 ± 0.97 vs. 7.00 ± 3.00, p = 0.12), $VIP_{Post}$ (0.81 ± 0.05 vs. 2.57 ± 0.26, $p = 3.1 \times 10^{-11}$) and $VIP_{Sup}$ (1.35 ± 0.11 vs. 0.76 ± 0.06, $p = 2.4 \times 10^{-6}$) neurons. **d.** No selection bias in peri-running analysis of Pyr neurons. Numbers of ROI imaged (42.1 ± 3.5) per FOV are not significantly different from those of fluorescently active ROI analysed (42.0 ± 3.5, p = 0.98) in Emx1-IRES-Cre;CaMK2a-tTA;Ai94 mice. On average 99.8 ± 0.2% of imaged Pyr neurons are included in data analysis. **e.** VIP activation significantly leads that of Pyr neurons in self-initiated running ($p = 1.6\times 10^{-18}$, Kolmogorov-Smirnov test). *** p<0.001, n.s.: not significant, Error bars: s.e.m.

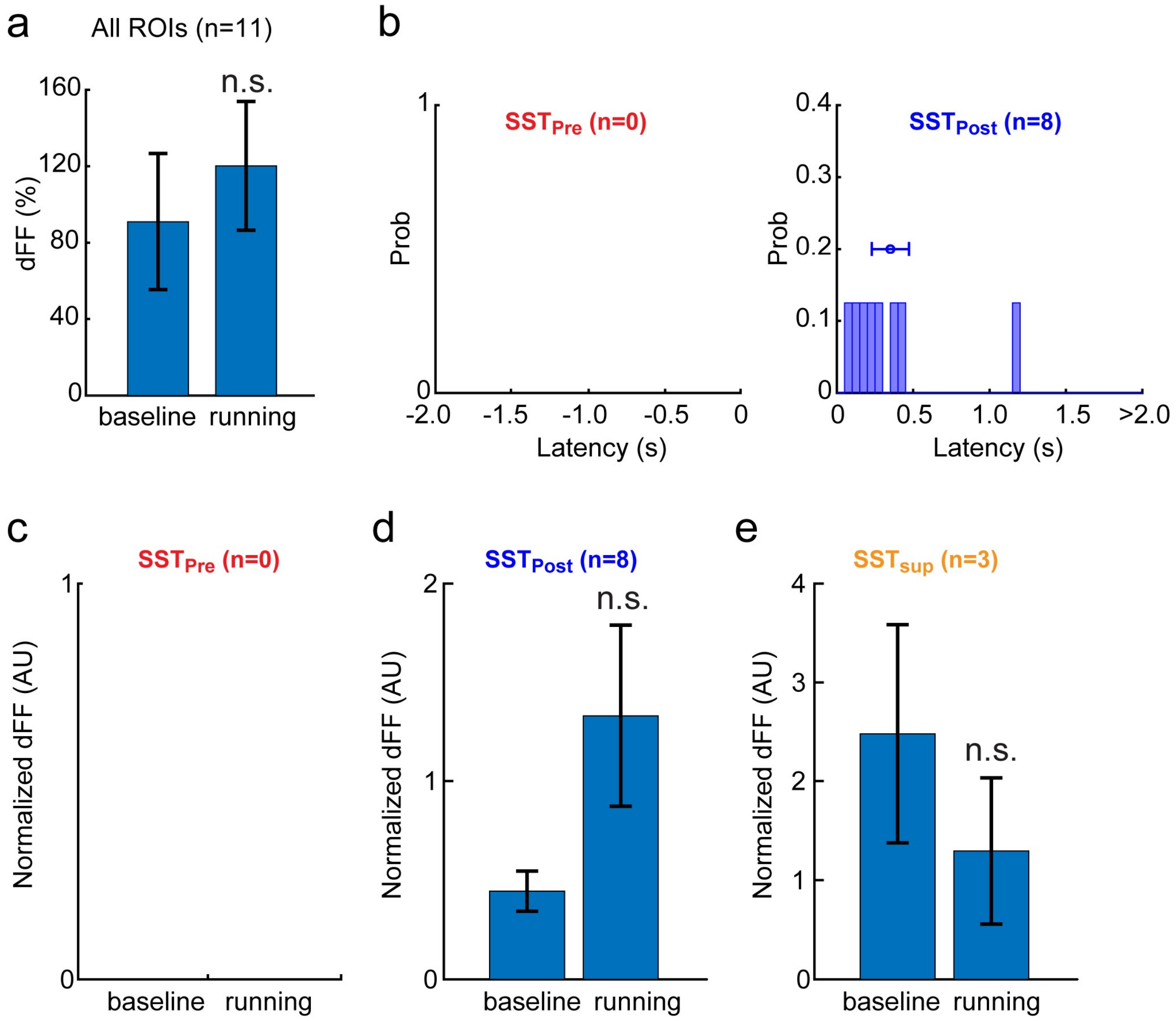


**Supplementary Figure 8. Extended data of example SST neurons in Fig. 3. a.** Mean ΔF/F activity of example SST neurons (n = 11) in Fig. 3a-c (baseline vs. running: 91.0 ± 35.7 vs. 120.2 ± 33.6, p = 0.56, *t*-test). **b.** Latency histograms of example $SST_{Post}$ neurons (0.350 ± 0.122 sec, right). No $SST_{Pre}$ cell was found in this FOV. **c - e.** Normalized ΔF/F activities during baseline imaging and running in example $SST_{Pre}$ (**c**), $SST_{Post}$ (**d**) and $SST_{Sup}$ neurons (**e**). Due to activity variations and limited sample size, running caused insignificant changes in $SST_{Post}$ (0.44 ± 0.10 vs. 1.33 ± 0.46, p = 0.08) and $SST_{Sup}$ (2.48 ± 1.10 vs. 1.29 ± 0.74, p = 0.42) neurons. No non-responsive SST neuron was found in this FOV. n.s.: not significant, Error bars: s.e.m.

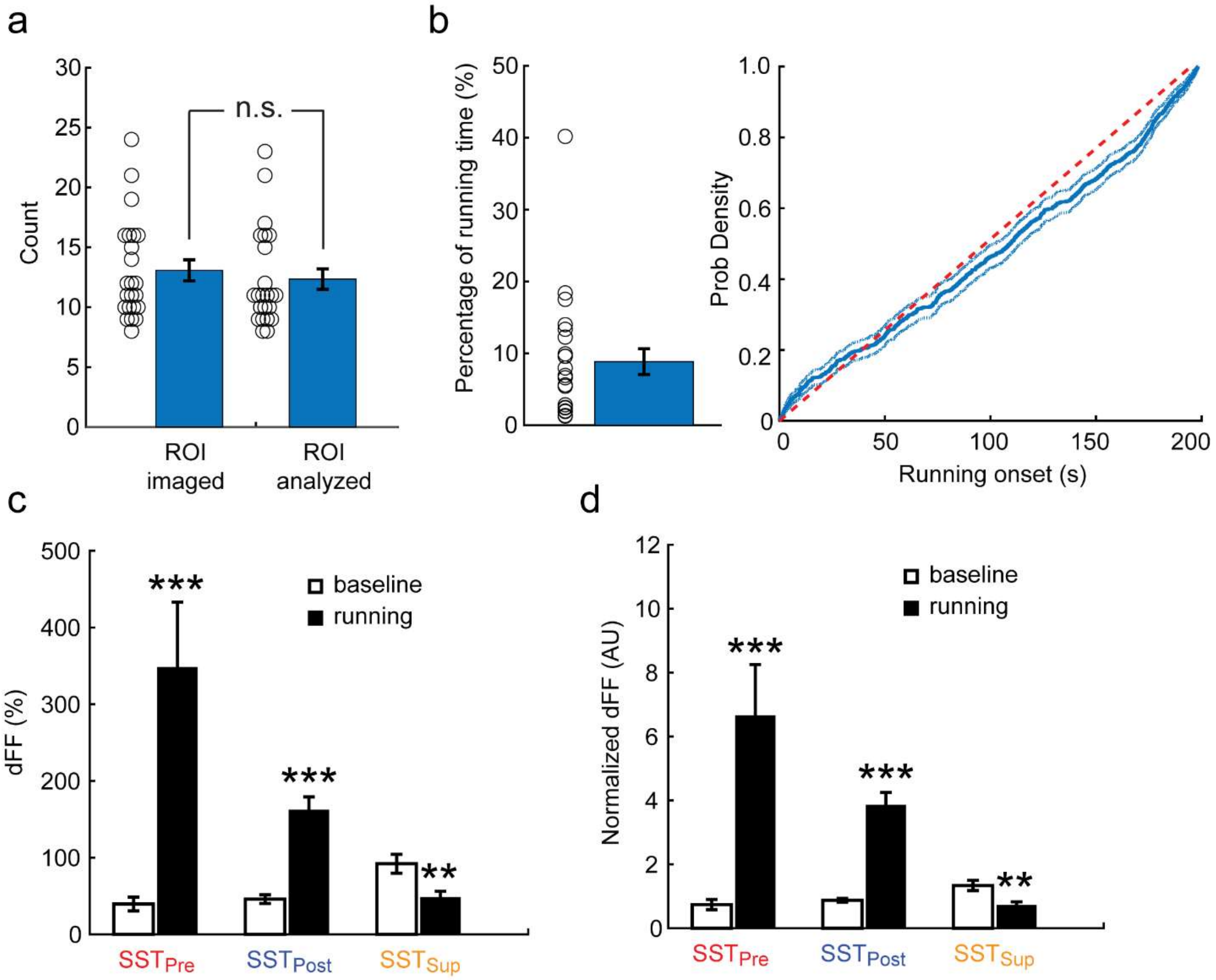


**Supplementary Figure 9. Population analysis of peri-running dynamics in SST neurons. a**. Numbers of ROI imaged (13.1 ± 0.9, mean ± s.e.m., same below, total 301) and fluorescently active ROI analyzed (12.3 ± 0.9, total 284, p = 0.5, *t*-test) per FOV in SST-IRES-Cre;Ai163 mice. Overall 94.6 ± 1.7% of imaged SST neurons are analyzed. **b**. Running statistics of SST-IRES-Cre;Ai163 mice. Left: percentage of running time (8.9% ± 1.8%) in 23 FOVs analyzed. Right: cumulative distribution of the running onset. **c**. Mean ΔF/F activities during baseline imaging and running in $SST_{Pre}$, $SST_{Post}$ and $SST_{Sup}$ neurons. At population level both $SST_{Pre}$ (39.54 ± 9.0 vs. 348.0 ± 85.0, n = 17, p = 0.001) and $SST_{Post}$ (45.9 ± 5.7 vs. 161.9 ± 17.3, n = 186, p = $5.7 \times 10^{-10}$) neurons significantly increase activities after running starts. However, $SST_{Sup}$ cells show significant activity suppression by running (92.1 ± 12.3 vs. 47.9 ± 8.3, n = 81, p = 0.0033). **d.** Consistent results from normalized ΔF/F activities during baseline and running in $SST_{Pre}$ (0.74 ± 0.16 vs. 6.65 ± 1.61, p = $9.1 \times 10^{-4}$), $SST_{Post}$ (0.88 ± 0.06 vs. 3.85 ± 0.40, p = $1.4 \times 10^{-12}$) and $SST_{Sup}$ (1.34 ± 0.16 vs. 0.71 ± 0.11, p = 0.0017) neurons. ** p<0.01, *** p<0.001, n.s.: not significant, Error bars: s.e.m.

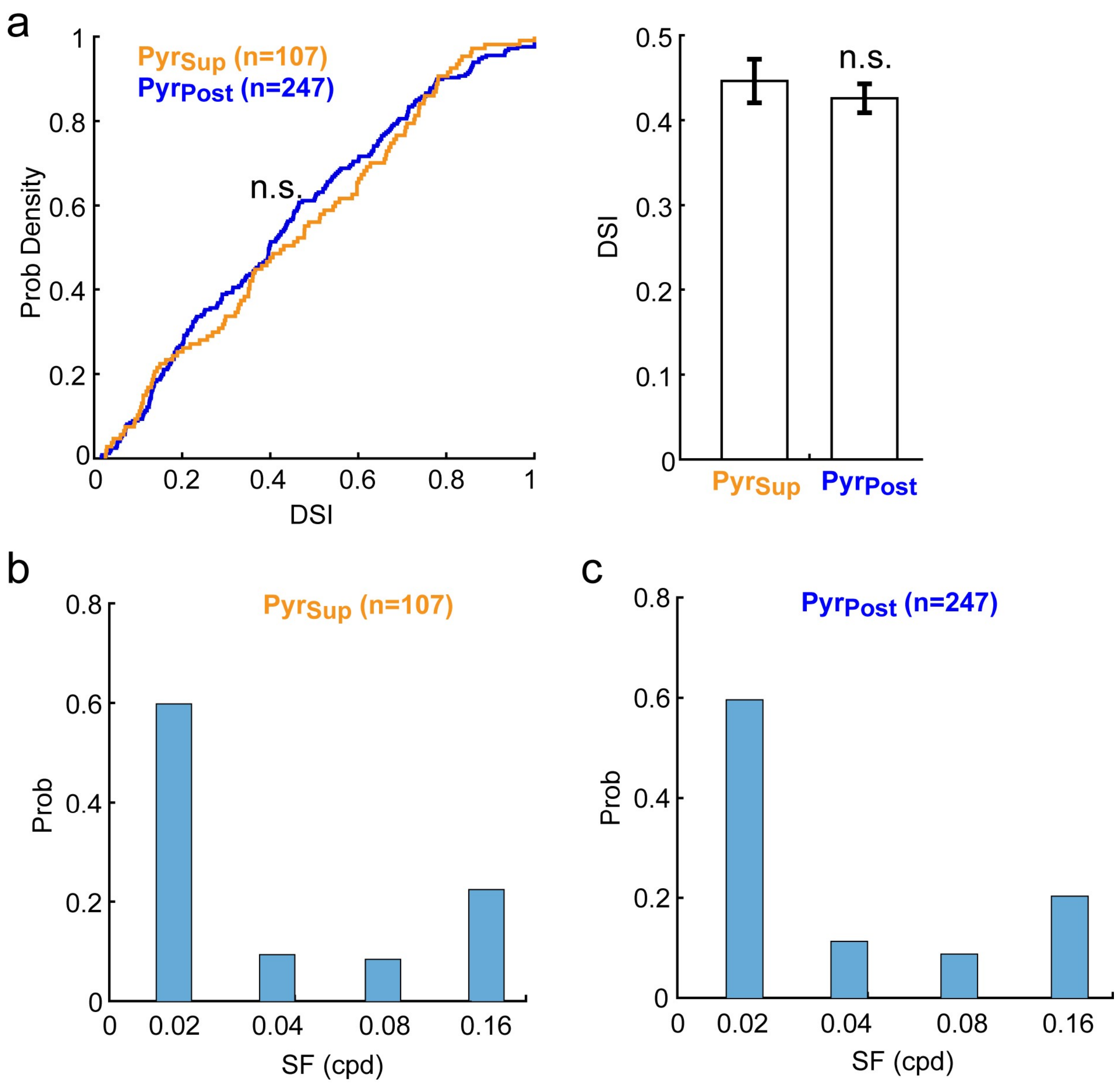


**Supplementary Figure 10. Response properties of Pyr neurons to drifting gratings. a**. No significant difference of DSI between $\text{Pyr}_{\text{sup}}$ and $\text{Pyr}_{\text{post}}$ neurons, as quantified by the cumulative distributions (left panel, p = 0.59, Kolmogorov-Smirnov test) and mean values (right panel, 0.446 ± 0.026 vs. 0.426 ± 0.017, p = 0.51, *t*-test). **b & c**. $\text{Pyr}_{\text{sup}}$ (**b**) and $\text{Pyr}_{\text{post}}$ (**c**) neurons show similar SF preferences. n.s.: not significant, Error bars: s.e.m.

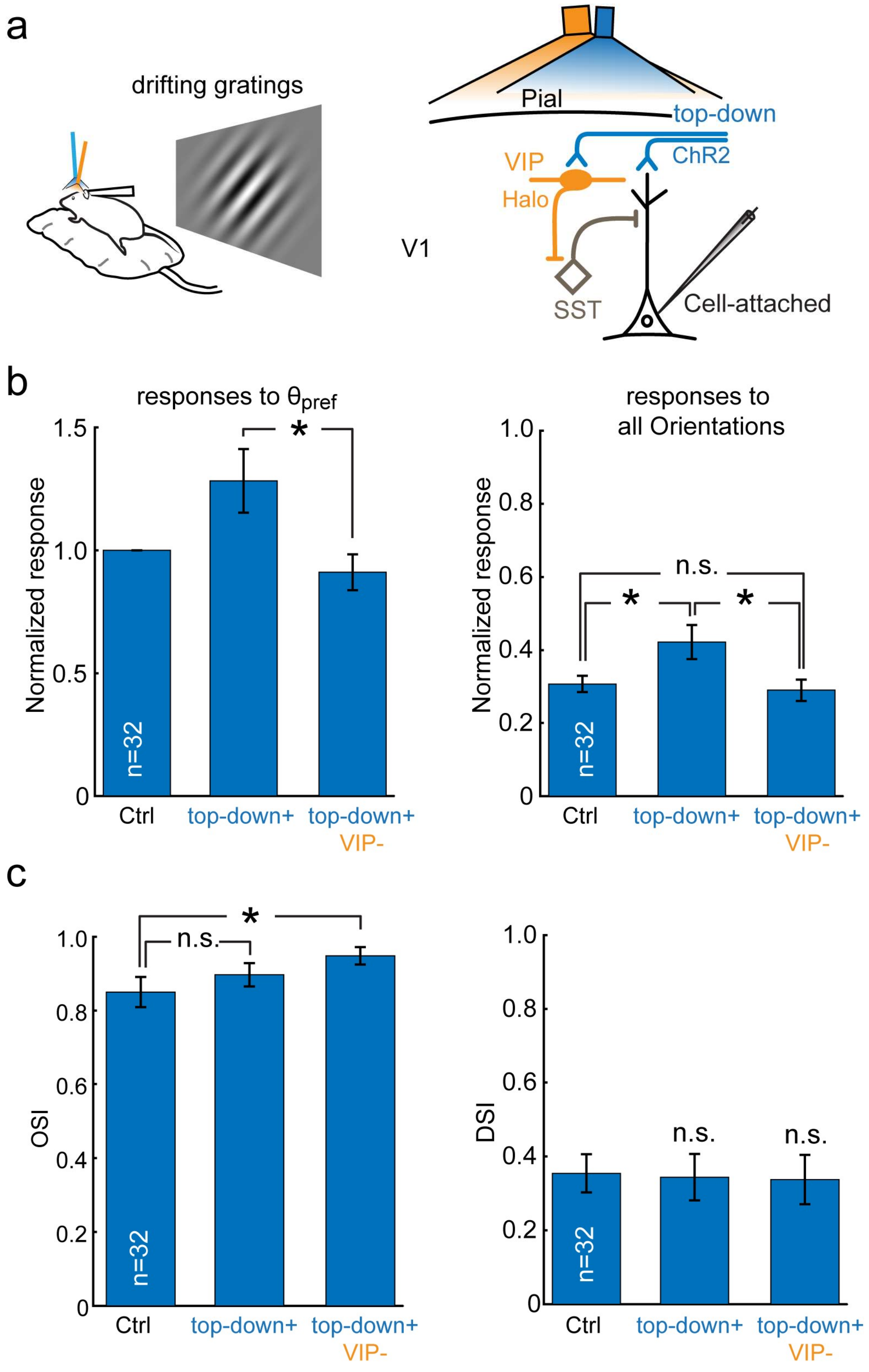

a
drifting gratings
Pial
top-down
VIP
ChR2
Halo
V1
SST
Cell-attached
b
responses to θpref
1.5
1.0
0.5
0
Normalized response
*
n=32
Ctrl
top-down+
top-down+
VIP-
responses to
all Orientations
1.0
0.8
0.6
0.4
0.2
0
Normalized response
n.s.
*
*
n=32
Ctrl
top-down+
top-down+
VIP-
c
1.0
0.8
0.6
0.4
0.2
0
OSI
*
n.s.
n=32
Ctrl
top-down+
top-down+
VIP-
1.0
0.8
0.6
0.4
0.2
0
DSI
n.s.
n.s.
n=32
Ctrl
top-down+
top-down+
VIP-

**Supplementary Figure 11. Optogenetic manipulation validates the VIP-SST-Pyr microcircuit mediates visual predictive processing. a**. Experimental setup. Cell-attached recordings of V1 Pyr neurons were made in anesthetised VIP-IRES-Cre;Ai39 mice (expressing halorhodopsin or Halo in VIP neurons, n = 9) previously receiving AAV-CaMKIIα-hChR2(H134R)-EYFP or AAV-DJ-CaMKII-hChR2(H134R)-EYFP injections in ACC-centred frontal regions. ChR2-expressing axons of frontal top-down projections were focally activated using a blue (wavelength 473 nm) laser coupled with optic fiber (200 μm in diameter) over V1. To inactivate Halo-expressing V1 VIP INs, an optic-fiber (600 μm in diameter) coupled yellow laser (593 nm) was used. Pyr responses to whole-screen drifting gratings were compared between no laser (ctrl), blue laser (top-down+) and blue+yellow lasers (top-down+/VIP-) conditions, representing control, top-down activation (resembling running) and inactivation of V1 VIP INs during running, respectively. **b.** Disinhibition of V1 Pyr neurons by focal activation of top-down axons in V1, which could be completely reversed by optical inactivation of VIP INs. Left panel: Normalized $\theta_{pref}$ response (top-down+ vs. top-down+/VIP-: 1.28 ± 0.13 vs. 0.91 ± 0.07, n = 32, p = 0.015, t-test). Right panel: average normalized response to all orientations (ctrl vs. top-down+: 0.31 ± 0.022 vs. 0.42 ± 0.044, n = 32, p = 0.03; top-down+ vs. top-down+/VIP-: 0.42 ± 0.044 vs. 0.29 ± 0.030, p = 0.019; ctrl vs. top-down+/VIP-: p = 0.64). **c**. Sharpening of the orientation tuning of V1 Pyr neurons. Left panel: Significant increase of Pyr OSI by optical inactivation of V1 VIP INs during top-down activation, but not by optical activation of top-down projections alone (ctrl vs. top-down+/VIP-: 0.850 ± 0.041 vs. 0.948 ± 0.024, n = 32, p = 0.04; ctrl vs. top-down+: 0.850 ± 0.041 vs. 0.897 ± 0.032, p = 0.38; top-down+ vs. top-down+/VIP-: p = 0.20). Right panel: Optogenetic manipulations have no effects on Pyr DSI (ctrl vs. top-down+/VIP-: 0.354 ± 0.052 vs. 0.337 ± 0.067, n = 32, p = 0.84; ctrl vs. top-down+: 0.354 ± 0.052 vs. 0.344 ± 0.063, n = 32, p = 0.90; top-down+ vs. top-down+/VIP-: p = 0.94). * p<0.05, n.s.: not significant, Error bars: s.e.m.
